# Genomic-Thermodynamic Phase Synchronization: Maxwell’s Demon-Like Regulation of Cell Fate Transition

**DOI:** 10.1101/2024.12.26.630379

**Authors:** Masa Tsuchiya, Kenichi Yoshikawa, Alessandro Giuliani

## Abstract

Dynamic criticality - the balance between order and chaos - is fundamental to genome regulation and cellular transitions. In this study, we investigate the distinct behaviors of gene expression dynamics in MCF-7 breast cancer cells under two stimuli: Heregulin (HRG), which promotes cell-fate transitions, and Epidermal Growth Factor (EGF), which binds to the same receptor but fails to induce cell-fate changes. We model the system as an open, non-equilibrium thermodynamic system and introduce a convergence-based framework for robust estimation of information-thermodynamic metrics.

Our analysis reveals that the Shannon entropy of the critical point (CP) dynamically synchronizes with the entropy of the rest of the whole expression system (WES), reflecting coordinated transitions between ordered and disordered phases. This phase synchronization is driven by net mutual information scaling with CP entropy dynamics, demonstrating how the CP governs genome-wide coherence. Furthermore, higher-order mutual information emerges as a defining feature of the nonlinear gene expression network, capturing collective effects beyond simple pairwise interactions.

By achieving thermodynamic phase synchronization, the CP orchestrates the entire expression system. Under HRG stimulation, the CP becomes active, functioning as a Maxwell’s demon with dynamic, rewritable chromatin memory to guide a critical transition in cell fate. In contrast, under EGF stimulation, the CP remains passive during non-critical expression dynamics.

These findings establish a biophysical framework for cell-fate determination, paving the way for innovative approaches in cancer research and stem cell therapy.

## Introduction

Dynamic criticality in gene expression, characterized by shifts in global expression profiles initiated by changes in specific gene subsets, plays a pivotal role in regulating genome dynamics and driving critical transitions [Liu et al., 2017]. This delicate balance between order and chaos enables genomic regulation to be both flexible and stable [Shmulevich et al., 2005; Zimatore et al., 2021].

Our previous studies demonstrated that this balance underlies both precise cellular responses to external signals and fundamental processes such as cell differentiation and embryonic development; these dynamics are central to understanding cell-fate changes. Our key findings are summarized below:

### 1. Emergent Self-Organization within a Genome Attractor

Self-organized critical (SOC) control of genome expression is quantified by the normalized root mean square fluctuation (*nrmsf*) of gene expression, which delineates distinct response domains (critical states). Large gene groups exhibit coherent stochastic behavior (CSB), converging around centers of mass (CM); in our approach, these CMs are treated as unit masses and act as attractors [Tsuchiya et al., 2014, 2015, 2016]. The whole-genome CM is the main genome attractor (GA), while local critical states serve as either subcritical (low-variance genes) or supercritical (high-variance genes) attractors. Cyclic fluxes among these attractors form an open thermodynamic “genome engine” that regulates genome expression [Tsuchiya et al., 2020, 2022].

### 2. CP as a Central Organizing Hub

A specific set of genes, termed the critical point (CP), exhibits bimodal, singular behavior based on the *nrmsf* metric. The CP acts as a central hub, spreading expression variability changes across the entire genome and driving critical transitions. When the system deviates from homeostasis, a state change in the CP propagates perturbations throughout the genome, demonstrating SOC in genomic regulation [Tsuchiya et al., 2023a].

### 3. Modeling the Genome Engine as a Dynamical System

The genome engine can be modeled as a one-dimensional dynamical system [Tsuchiya et al., 2023b]. In this model, the CM’s expression level at time ε_t_ represents “position,” its first difference (ε_t+1_ – ε_t_) represents “momentum,” and its second difference (ε_t+1_ – 2ε_t_ + ε_t-1_) acts as an “effective force” driving energy changes. This framework, grounded in stochastic thermodynamics [Parrondo et al., 2015; Ito, 2018; Peliti and Pigolotti, 2021; Shiraishi, 2023; Korbel and Wolpert, 2024], illustrates how the genome maintains a near-balance between influx and outflux while dynamically interacting with its environment.

### 4. Transition-Driven Switching of Cyclic Flows

Critical transitions in the genome engine reverse cyclic expression fluxes [Tsuchiya et al., 2020, 2022]. These transitions are linked to structural changes, such as the bursting of peri-centromeric-associated domains (PADs) in MCF-7 cancer cells, affecting chromatin folding and enabling dynamic genomic regulation [Zimatore et al., 2021; Krigerts et al., 2021; Erenpreisa et al., 2023].

### 5. OMG-CP-GA Network Synchronizing CP and GA for Cell-Fate Change

The OMG-CP-GA network governs cell-fate transitions by coordinating interactions among oscillating-mode genes (OMGs), the CP, and the GA. OMGs, identified by high scores on the second principal component (PC2, with PC1 representing the equilibrium gene expression profile) [Zimatore et al., 2021], modulate synchronization between the CP and GA. This synchronization maintains genome-wide balance and triggers critical transitions, leading to coordinated shifts in genome expression and chromatin remodeling [Tsuchiya et al., 2023a].

### 6. Advancements over Classical Self-Organized Criticality Models

Unlike classical SOC (cSOC) models, which involve state transitions from subcritical to supercritical toward a critical attractor [Bak et al., 1987, 1988; Bak and Chen, 1991; Watkins et al., 2016], our SOC model features a dynamic CP that actively induces state changes to guide cell-fate transitions. This mechanism enables the genome to adaptively regulate itself in response to stimuli [Tsuchiya et al., 2016, 2020, 2022].

### 7. Universality Across Biological Systems

The SOC control of genome expression has been demonstrated in distinct biological systems, including cell differentiation and embryonic development: HRG- and EGF-stimulated MCF-7 human breast cancer cells [Saeki et al., 2009]; atRA- and DMSO-stimulated HL-60 human leukemia cells [Huang et al., 2005]; Th17 cell differentiation [Ciofani et al., 2012]; early embryonic development in mice [Deng et al. 2014] and human embryonic development [Yang et al., 2013]. These findings highlight the robustness and universality of the dynamic criticality model [Giuliani et al., 2017; Tsuchiya et al., 2016, 2020, 2022].

Building on these findings and utilizing our dynamic approach, this study aims to:

1. Elucidate how non-equilibrium thermodynamics of open systems governs genome-wide expression;
2. Distinguish between effective and non-effective dynamics in cell-fate transitions;
3. Propose a unified framework that integrates expression flux analysis with genomic thermodynamics.

We analyze temporal gene expression data from MCF-7 cancer cell lines stimulated with epidermal growth factor (EGF), which promotes proliferation without altering cell fate, and heregulin (HRG), which induces commitment to differentiation following rapid dissipation of an inital gene expression perturbation [Saeki et al., 2009; Zimatore et al., 2021; Erenpreisa et al., 2023]. By modeling these two processes as open stochastic thermodynamic systems - characterized by exchanges of heat and matter (e.g., gases, nutrients, and waste products) with their surroundings and by inherent randomness-we capture the dynamic and non-deterministic nature of genome regulation influenced by both external and internal factors.

To assess differences in genome expression dynamics, we use Shannon entropy and mutual information as quantitative measures related to disorder and predictability. Shannon entropy, rooted in information theory, quantifies unpredictability in biological processes and relates to entropy production in non-equilibrium systems, linking informational disorder to energy dissipation. This framework helps explain how biological systems maintain order and perform work under stochastic, far-from-equilibrium conditions.

In contrast, mutual information captures interdependencies, feedback, control mechanisms, thermodynamic efficiency, and information flow [Shannon, 1948; Adami, 2004; Sagawa and Ueda, 2012a, 2012b; Barato and Seifert, 2014]. A foundational perspective on the interplay between information and thermodynamics is illustrated by Maxwell’s demon, a thought experiment introduced by James Clerk Maxwell [Maxwell, 1871]. Although the demon appears to reduce entropy, thereby violating the second law of thermodynamics, extensive studies [Szilárd, 1929; Landauer, 1961; Astumian, 1997; Bennett, 2003; Toyabe et al., 2010; Bérut et al., 2012; Sagawa and Ueda, 2012a, 2012b; Seifert, 2012; Flatt et al., 2023; Annby-Andersson et al., 2024] have shown that information processing entails a thermodynamic cost, preserving the second law.

Toyabe et al. (2010) first experimentally demonstrated the conversion of information into usable work via real-time feedback control, effectively realizing a modern Maxwell’s demon using a colloidal particle system. Building on this, Sagawa and Ueda (2012) and Parrondo et al. (2015) formalized a general thermodynamic framework in which the demon operates as a memory through three sequential phases: initialization (or preparation), measurement, and feedback, each carrying a well-defined energetic cost. Subsequent non-quantum experiments have continued to validate and extend this paradigm [Koski et al., 2014; Chida et al., 2017; Ribezzi-Crivellari and Ritort, 2019].

Far from being merely a metaphor, Maxwell’s demon embodies a functional principle, underscoring the profound connection between information processing and thermodynamic behavior in non-equilibrium open systems, including within biological phenomena.

### The overview and objectives of our study are as follows

Understanding complex gene expression networks requires accurate estimation of information-theoretic metrics such as Shannon entropy and mutual information, which quantify disorder, predictability, and interdependencies. These metrics are challenging to estimate due to the stochastic nature of gene expression and the complexity of the underlying networks.

To address this challenge, we exploit coherent stochastic behavior (CSB) through a convergence-based approach [Tsuchiya et al., 2014, 2015, 2016]. This enhances the estimation of the critical point (CP) location along the axis of gene expression variability (**section 2.1**) and ensures robust identification of CP genes by grouping them on the basis of their expression variability (**section 2.2**). Additionally, this approach improves the robustness of information-thermodynamic metrics, including entropy and mutual information, which quantify the dynamics of genomic regulation (**section 2.3**). This methodology allows us to characterize the genomic mechanisms distinguishing stimuli that do not promote cell-fate change (EGF stimulation) from those that drive cell-fate transitions (HRG stimulation).

Our study focuses on the following objectives:

1. **Open Thermodynamic Modeling and Phase Synchronization** (**section 2.4**): Develop a data-driven approach to model an open stochastic thermodynamic system of MCF-7 cell lines and investigate how order-disorder phase synchronization between the CP and the whole expression system (WES) regulates genome expression under EGF and HRG stimulation.
2. **Higher-Order Mutual Information** (**section 2.4**): Investigate complex nonlinear interdependencies in networks within the WES by identifying higher-order mutual information.
3. **CP as Maxwell’s Demon-Like Rewritable Chromatin Memory** (**sections 2.5** and **2.6**): Examine the CP’s role as a Maxwell’s demon-like rewritable chromatin memory, clarify its function in entropy regulation and information flow, and thereby deepen our understanding of genomic thermodynamics.
4. **Gene Expression Variability as a Proxy for Chromatin Remodeling Dynamics (section 2.7):** Investigate whether gene expression variability serves as a proxy for underlying chromatin remodeling dynamics by analyzing experimental evidence.

In the Discussion section, we expand on our findings and explore their potential implications for controlling the dynamics of cancer cell fate through three key perspectives: (1) the order parameter of phase synchronization as a bridge between dynamical systems and non-equilibrium thermodynamics; (2) autonomous genome computing as a conceptual framework; and (3) future prospects for genome intelligence (GI). Finally, the main insights are summarized in the Conclusion section.

## 2. Results

### 2.1 Normalized Temporal Variability of Gene Expression as a Metric of Self-Organization and a Proxy for Chromatin Flexibility

To identify the critical point (CP) genes, which exhibit bimodal singular behavior, we introduce a metric parameter that quantifies the self-organization of time-series whole-genome expression data obtained from both microarray and RNA-Seq datasets (see methodological details in [Tsuchiya et al., 2023b]). This metric parameter is defined by the root mean square fluctuation (*rmsf*_i_), representing the standard deviation of a gene’s expression levels over time, calculated as follows:

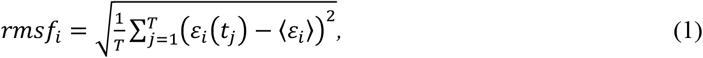

where *ε*_*i*_(*t*_*j*_) (*i*= 1, 2, ‥, *N*) is the expression level of the *i*^th^ gene at a specific cell state or experimental time point *t*_*j*_, ⟨*ε*_*i*_⟩ is the average expression level of the *i*^th^ gene over all time points; and *T* is the total number of cell states or experimental time points.

To compare gene expression variability across different biological systems [Tsuchiya et al., 2020, 2022], we normalize the *rmsf*_i_ value by the maximum *rmsf* observed in the dataset, resulting in the normalized *rmsf* (*nrmsf*_i_). For each gene expression value measured across experimental time points, we first compute *nrmsf*_i_ and then take its natural logarithm, *ln*(*nrmsf*_*i*_) as well as the natural logarithm of gene expression value, *ln*(*ε*_*i*_(*t*_*j*_)). In our study, “expression” specifically refers to this log-transformed value. This transformation captures scaling response behaviors in SOC control [Tsuchiya et al., 2015, 2016, 2023b; Giuliani et al., 2017] and reduces noise.

For microarray data, which rarely contain zero values, this approach works effectively. In contrast, RNA-Seq data, often characterized by numerous zero values, require the addition of small random noise to zero entries, followed by ensemble averaging and then logarithmic transformation, in order to preserve coherent stochastic behavior (CSB) [Tsuchiya et al., 2023b].

The use of *ln*(*nrmsf*) as a ranking parameter is biologically justified because gene expression variability, as quantified by *ln*(*nrmsf*), reflects chromatin flexibility. Greater variability indicates that chromatin is more accessible and dynamically regulated. In HRG-stimulated MCF-7 cancer cells undergoing a cell-fate transition, principal component analysis (PCA) reveals that principal component scores accurately predict each gene’s *ln*(*nrmsf*). In particular, the second and third principal components (PC2 and PC3) capture chromatin structural flexibility associated with chromatin remodeling during the critical transition that guides cell fate [Zimatore et al., 2021]. Additional experimental evidence presented in **section 2.7** further supports the use of *ln*(*nrmsf*) as a proxy for chromatin flexibility [Krigerts et al., 2021; Erenpreisa et al., 2023].

The CP genes, identified by a critical range of *ln*(*nrmsf*) values, serve as regulatory hubs that orchestrate genome-wide transitions [Tsuchiya et al., 2020, 2022, 2023a]. In this study, we demonstrate that activation of CP genes at a critical transition drives coherent chromatin remodeling and explore how this process parallels Maxwell’s demon–like behavior (see **sections 2.5** and **2.6**).

### 2.2 Identifying CP Genes Using the Metric Parameter

The whole-genome mRNA expression dataset is analyzed using *ln*(*nrmsf*) as the metric. Genes are first sorted in descending order of *ln*(*nrmsf*) and then grouped into clusters large enough to capture genome-wide averaging behavior of collective expression dynamics [Tsuchiya et al., 2014, 2015, 2016]. CSB in gene expression emerges when the sample size (*n*) exceeds 50 [Censi et al., 2011]. As the sample size increases, the group’s center of mass (CM) with unit mass converges to a constant value, driven by the law of large numbers. This convergence reflects a nearly balanced exchange between the inflow and outflow of expression flux within the whole expression system (WES) [Tsuchiya et al., 2023b] (see more in **section 2.4**). To capture such types of thermodynamic behavior, each group needs to contain more than 200 genes.

**Figure 1** illustrates the data-processing workflow for time-series whole gene expression, which underpins the information-thermodynamic analysis (ITA) and the computation of the associated information-thermodynamic metrics. This framework establishes a data-driven approach for modeling the gene expression dynamics of MCF-7 cells as an open stochastic thermodynamic system (see **section 2.3**).

**Figure 1.**
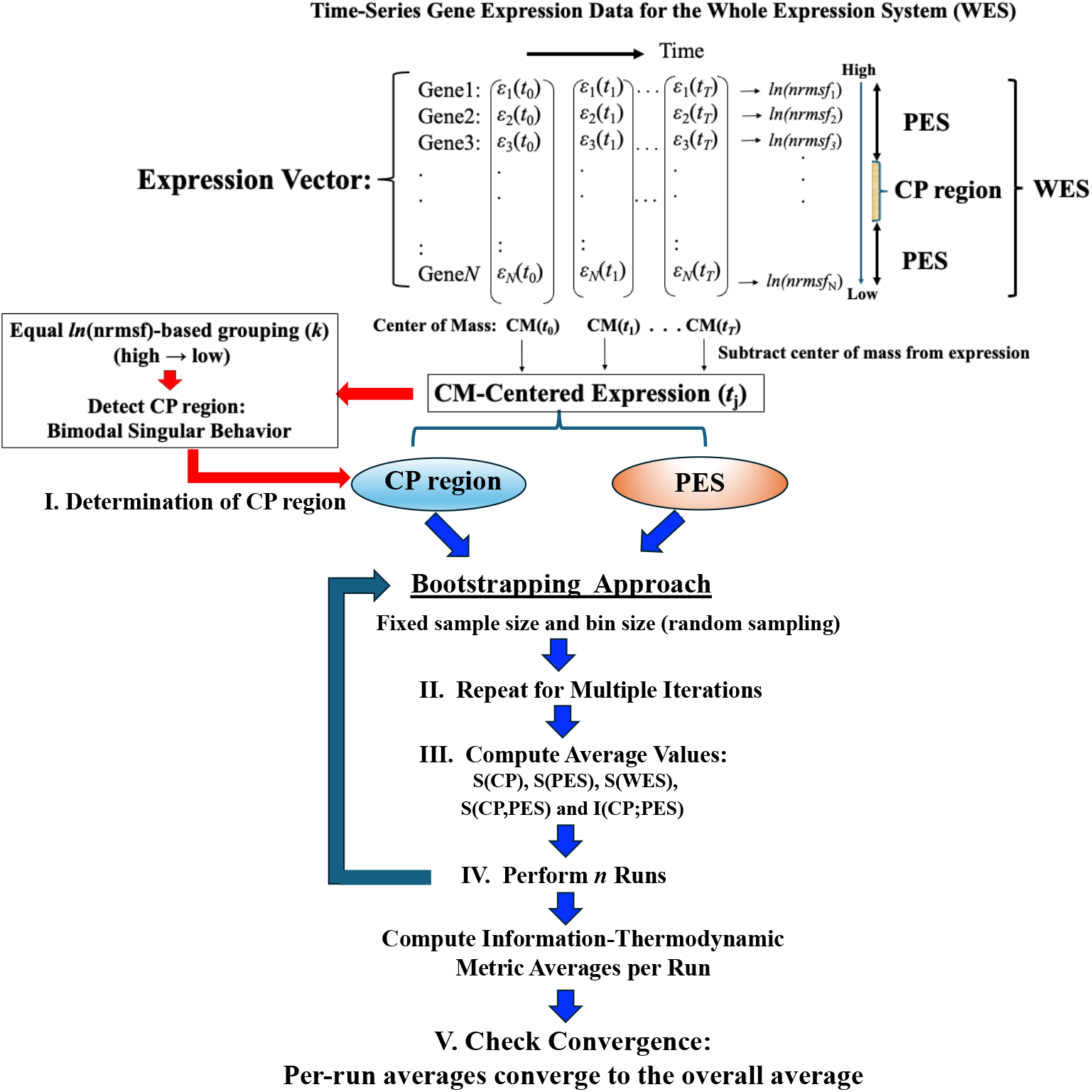
Workflow for Computing Information-Thermodynamics Metrics from Time-Series Gene Expression Data. This figure illustrates a five-step workflow for assessing information-thermodynamics metrics from time-series gene expression data of MCF-7 cells. The process begins by constructing an *N*-dimensional expression vector (*N* = 22,277, corresponding to the number of genes on the array) for the whole expression system (WES) at each time point using normalized raw expression data (see **Material**). For each gene, the *nrmsf* value is then computed and transformed using the natural logarithm, along with the natural logarithm of the gene expression values (see **section 2.1** for details). The log-transformed gene expression values, after subtraction of the time-dependent center of mass (CM), are then sorted in descending order based on their *ln*(*nrmsf*) values. From this sorted data: **I**. The critical point (CP) region is determined by identifying a specific set of gene expressions that exhibit bimodal singular behavior, using equal-sized *ln*(*nrmsf*)-based grouping (*k* = 40 groups; see **Figure 2**). To estimate robust CP entropy, the CP region, defined as the range of *ln*(*nrmsf*) values exhibiting bimodal singular behavior, is standardized by adjusting its original interval by 0.2 units (see **Figure 2**). This adjusted range includes both CP genes and adjacent edge genes influenced by CP dynamics, thereby providing a sufficient gene set. **II**. The peripheral expression system (PES) is defined as the remaining portion of the WES that interacts with the CP region (**Figure 3**). **III**. Based on the CP and PES expression vectors, bootstrapping is performed using fixed parameters (random sample size = 1,000; bin size = 30), while varying the number of bootstrap iterations (see **Figure 4**), and the average values of the resulting metrics are computed (see details in **section 2.3**). **IV**. The entire bootstrapping process (run) is repeated *n* times (*n*= 10 in this study), with averaged metric values computed for each individual run. **V**. Convergence is confirmed when the average absolute difference between successive bootstrap iterations (500 and 1000 iterations for mutual information; 200 and 500 iterations for entropy) across all time points falls below 10^−3^. Definitions of the computed metrics (e.g., *S*(CP), *I*(CP;PES)) are provided in the main text.

The objective is to investigate how phase synchronization between the critical point (CP) and the whole expression system (WES) regulates genome-wide expression in response to EGF and HRG stimulation. A particular focus is on identifying distinct information-thermodynamic differences between conditions that induce cell-fate change (HRG) and those that do not (EGF), despite both ligands signaling through the same receptor.

As illustrated in **Figure 2**, the CP region is identified through temporal changes in overall expression along the *ln*(*nrmsf*) metric; genes within this region exhibit distinct bimodal singular temporal behavior. This characteristic behavior serves as a basis for distinguishing the peripheral expression system (PES) from the whole expression system (WES). In HRG-stimulated MCF-7 cells, the CP region and PES comprise 3,846 and 18,431 genes, respectively, while in EGF-stimulated MCF-7 cells, they comprise 4,033 and 18,244 genes, respectively. Hereafter, the CP region is referred to simply as the CP genes, unless a distinction between the region and the specific gene set is necessary.

**Figure 2.**
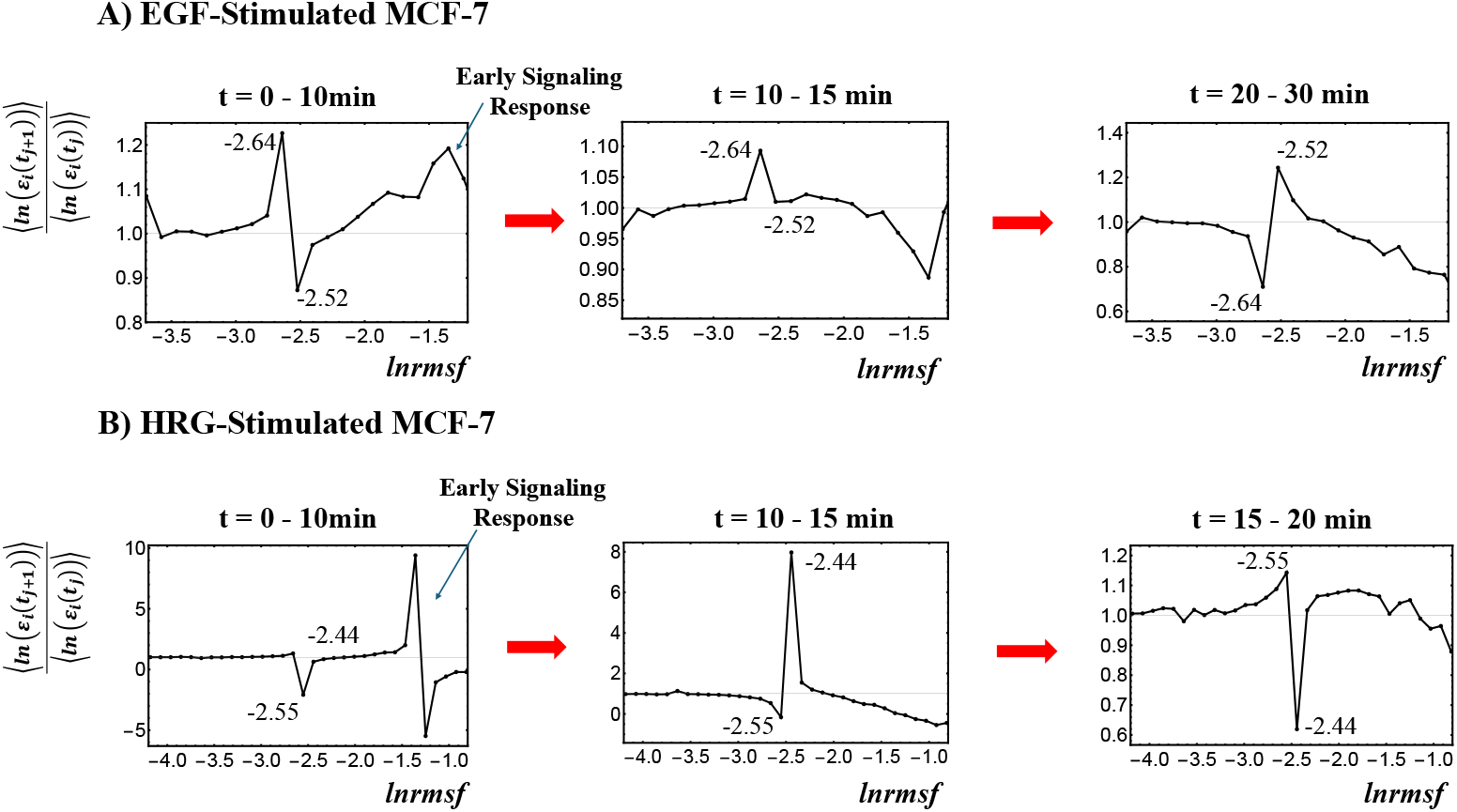
Identification of the Critical Point (CP) Region with Distinct Response Domains. **A)** EGF-stimulated MCF-7 cells and **B)** HRG-stimulated MCF-7 cells. This figure details step (I) from the workflow presented in **Figure 1**. The entire gene expression dataset is sorted and divided into 40 equal-sized groups based on *ln*(*nrmsf*) values, excluding groups containing fewer than 200 genes due to limited convergence, which typically occurs at the extreme high and low ends of the distribution. Because *ln*(*nrmsf*) values are time-independent, they are plotted on the *x*-axis. The y-axis represents the ratio of group ensemble averages across different time points, in which the CP region distinctly emerges with bimodal singular behavior. A black solid dot represents the natural logarithm of the ensemble average of the *nrmsf* value, <*nrmsf*_*i*_> for the *i*^*th*^ group, while the *y*-axis shows the ratio of the group’s average expression between consecutive time points, <*ln*(*ε*_i_(*t*_j+1_))>/<*ln*(*ε*_i_(*t*_j_))>. Under EGF stimulation, CP genes exhibit bimodal critical behavior with peaks ranging from –2.64 to –2.52 *ln*(*nrmsf*), corresponding to a CP region of –2.7 < *ln*(*nrmsf*) < –2.5 (4,033 genes). Under HRG stimulation, the peaks range from –2.55 to –2.44 *ln*(*nrmsf*), with a CP region of –2.6 < *ln*(*nrmsf*) < –2.4 (3846 genes). The CP region effectively separates high-variance from low-variance responses, thereby highlighting distinct regulatory domains within the genome. For details regarding the early signaling response, refer to section 2.4.

### 2.3 Open Stochastic Thermodynamic Model of MCF-7 Cell Line

This section presents a stochastic thermodynamic model of MCF-7 breast cancer cell line, focusing on the interaction between the critical point (CP) genes and the rest of the whole gene expression system (WES). The remaining genes in WES are collectively referred to as the Peripheral Expression System (PES).

We adopt an open thermodynamic framework in which cells continuously exchange heat and substances (gases, nutrients, and waste products) with their environment. This exchange influences gene expression dynamics. By analyzing temporal variations in entropy within the system, we gain insights into how entropy constrains the organization and stability of genomic regulation. This approach underscores the principles of non-equilibrium thermodynamics in open systems that govern genome-level interactions.

As illustrated in **Figure 3**, both the CP genes and the PES exchange energy with the external environment (OUT). We analyze an ensemble of CP genes and the PES within the cancer cell population, examining their entropy and interactions to uncover the open thermodynamic principles governing the temporal evolution of the system.

**Figure 3.**
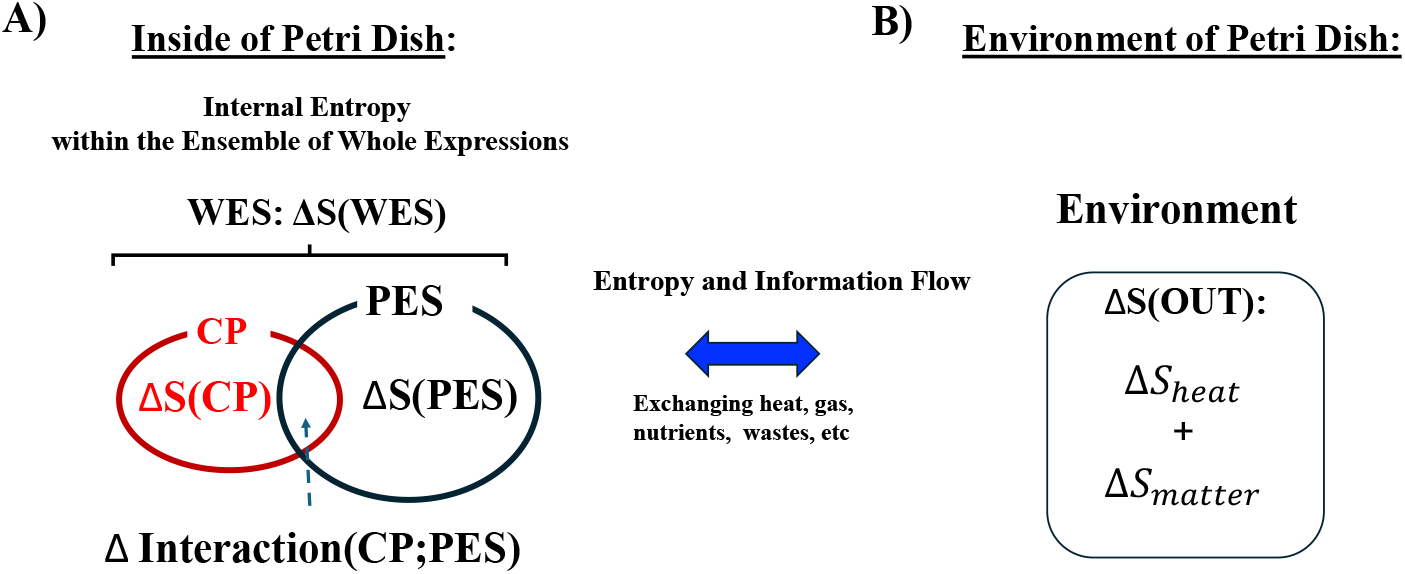
Open Stochastic Thermodynamic Model Setup for MCF-7 Cell Lines. This schematic depicts an open thermodynamic system for MCF-7 cell lines, where heat, gases, nutrients, and wastes are continuously exchanged with the environment. The system includes the whole expression system (WES), which comprises the critical point (CP) genes and the remaining genes, collectively referred to as the peripheral expression system (PES). Each component has its own entropy: *S*(WES), *S*(CP), and *S*(PES). **A)** Within the system, the total entropy change of WES (Δ*S*(WES)) is decomposed into changes in the CP (Δ*S*(CP)), the PES (Δ*S*(PES)), and their interaction term (ΔInteraction(CP;PES)). This decomposition enables the investigation of how these interaction terms relate to the mutual information Δ*I*(CP;PES). **B)** Entropy exchange with the environment (Δ*S*(OUT)) is composed of contributions from heat (Δ*S*_heat_) and matter exchange (Δ*S*_matter_). We apply the CSB method to estimate entropy and mutual information, demonstrating consistent convergence behavior (see **Figure 4**).

**Figure 4** illustrates that, with a fixed sample size (30), increasing the number of sampling repetitions via bootstrapping enables robust estimation of information-thermodynamic metrics. This approach leverages CSB to achieve convergence of average values across samplings, thereby stabilizing the estimates and revealing the thermodynamic principles governing the genome’s dynamic regulatory mechanisms. Additionally, the second thermodynamic condition is considered, accounting for entropy flow between the cell line and its external environment (beyond the confinement of the Petri dish), as described below.

**Figure 4.**
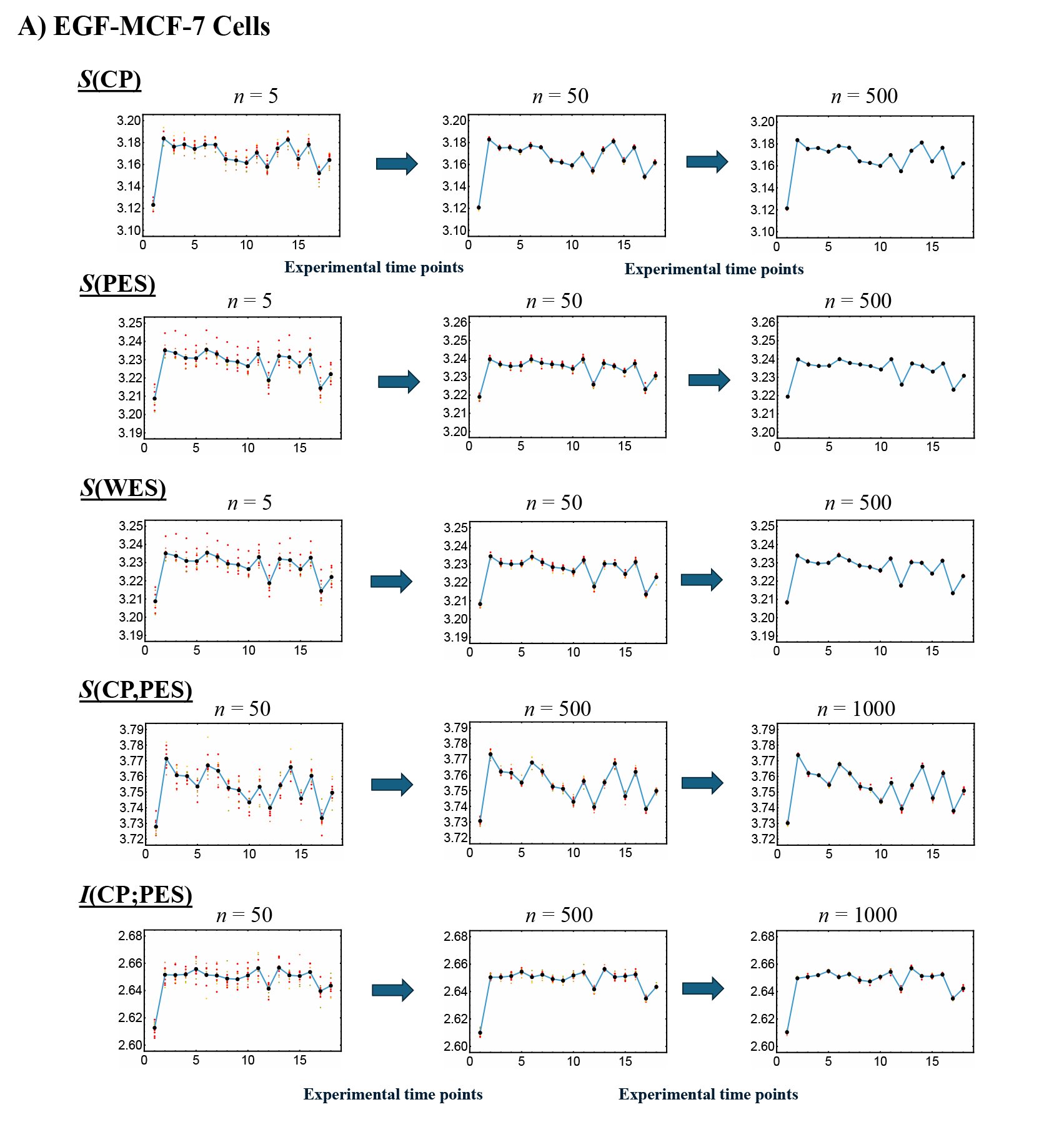
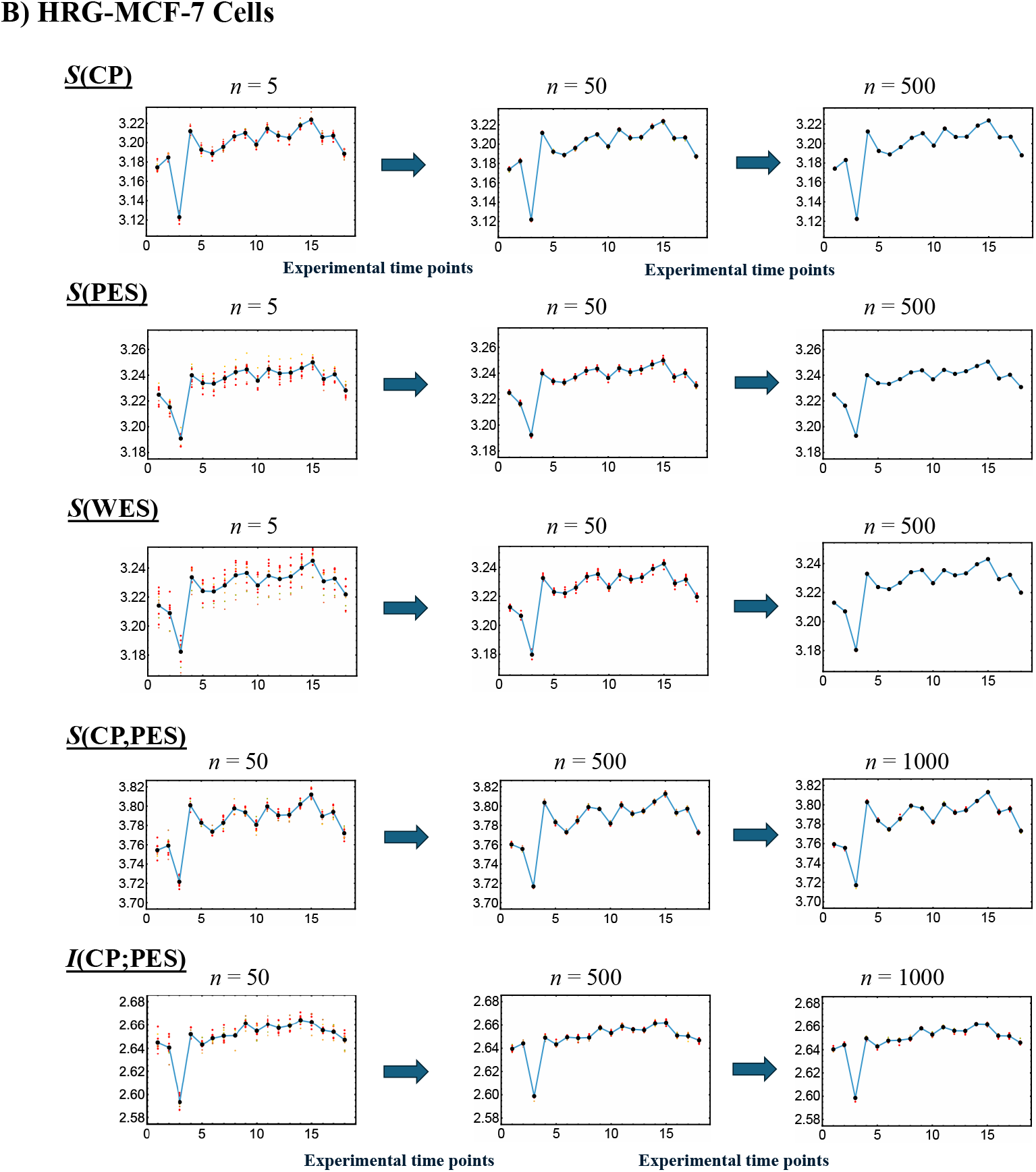
Convergence-Based Approach for Robust Estimation of Information-Thermodynamic Metrics. Panels **A)** EGF-stimulated and **B)** HRG-stimulated MCF-7 cells. This figure illustrates the details of step V) in the workflow outlined in **Figure 1**, focusing on convergence assessment. The approach enables robust estimation of key information-thermodynamic metrics: entropy of the CP (*S*(CP)), the WES (*S*(WES)), and the PES (*S*(PES)); as well as joint entropy (*S*(CP,PES)) and mutual information (*I*(CP;PES)). Each row shows the progression of convergence for a specific metric, with estimates computed via bootstrapping (sample size = 1,000; bin size = 30), following the square root rule of the sample size as described in [Tsuchiya et al., 2010]. The number of iterations increases across columns to illustrate convergence dynamics: entropy metrics are computed using 5, 50, and 500 iterations, while joint entropy and mutual information use 50, 500, and 1,000 iterations due to their higher dimensionality. To assess convergence, the entire procedure (run) is repeated 10 times under each condition, and the resulting metric values at each time point are plotted. Convergence is visually evaluated by the reduction in scatter among the 10 averaged runs, as highlighted by red solid circles approaching the overall average (black solid circles). Joint entropy and mutual information typically exhibit slower convergence because they require estimating two-dimensional joint frequency distributions, unlike the one-dimensional distributions used for entropy. The *x*-axes represent experimental time points, while the *y*-axes denote the corresponding metric values.

#### 1: Random Sampling

We repeatedly perform bootstrap random sampling on genes from the CP, PES and WES datasets, allowing for replacement (bootstrap sampling).

#### 2: Calculating Entropy

For each sample, we calculate the Shannon entropy using the following equation:

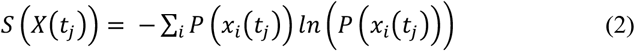

where *S*(X(*t*_j_)) is dimensionless; *X*(*t*_j_) represents CP(*t*_j_), PES(*t*_j_) or WES(*t*_j_) and *P*(*x*_i_(*t*_j_)) is the probability of gene *x*_i_ being expressed at *t*_j_.

Note that regarding **Figure 4**, we observed that varying the bin sizes used to calculate the probability distributions leads to consistent offsets in the resulting information-thermodynamic metrics (e.g., entropy), with larger bin sizes producing higher values. This behavior is exactly as expected for entropy as a proper state function. We chose a bin size of 30 for the probability distributions, following the square root rule (bin size 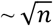, where *n* is the sample size), as described in [Tsuchiya et al., 2010].

#### 3: Relating Entropies and Higher-Order MI (Mutual Information)

We compute the joint entropy *S*(CP, PES) at *t*_j_:

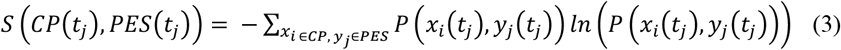

where *P*(*x*_i_(*t*_j_), *y*_j_(*t*_j_)) is the joint probability of CP gene *x*_i_ and PES gene *y*_j_ being simultaneously expressed at *t*_j_. To estimate the joint probability distribution between CP and PES genes, we construct a two-dimensional joint frequency table at each time point using a bootstrapping approach and a fixed bin size of 30.

The standard MI between CP and PES, *I*(CP: PES) is given by

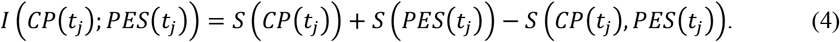

The joint entropy *S*(CP, PES) can also be expressed as internal entropy:

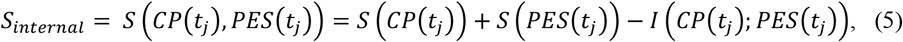

where the mutual information is subtracted to remove shared information, leaving only the unique and combined uncertainties of the two components. From this point onward, the time index *t*_j_ will be omitted for simplicity, except when explicitly necessary.

Note that WES consists of the CP genes and the remaining WES genes (PES). The entropies *S*(WES), *S*(CP), and *S*(PES) are calculated from their respective gene sets using the same bootstrap procedure and a dimensionless entropy measure, accounting for interactions with the external environment (**Figure 3**). As shown in **Figure 5**, *S*(WES) differs from *S*_internal_ (equation (5)) at all time points, indicating that the standard MI formula (equation (4)) does not fully capture all dependencies in MCF-7 cancer cells:

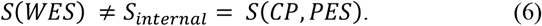

**Figure 5.**
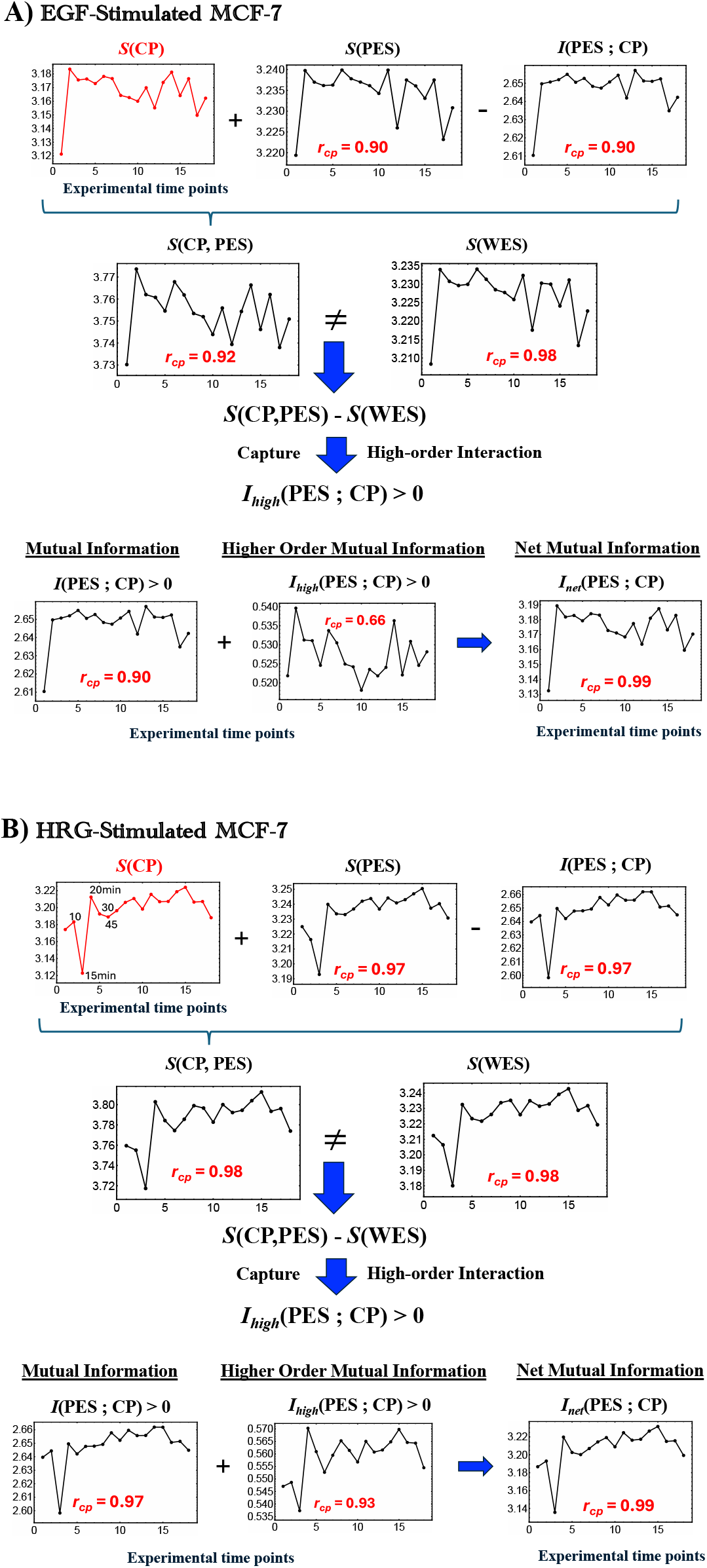
Thermodynamic Phase Synchronization and Higher-Order MI in MCF-7 Cells. **A)** EGF stimulation; **B)** HRG stimulation. Using a bootstrapping approach (500 iterations for entropy and 1000 iterations for MI and joint entropy, as described in **Figure 4**), we calculated the entropies *S*(WES), *S*(CP), and *S*(PES). These figures highlight the following three key points: 1) According to the standard MI framework, the relation *S*(WES) = *S*(CP) + *S*(PES) – *I*(CP; PES), where *S*(WES) = *S*(CP, PES) is expected to hold. However, the observed non-zero difference between *S*(WES) and the joint entropy *S*(CP, PES) indicates additional positive higher-order MI, *I*_*high*_(CP; PES)> 0. This implies that the net MI, *I*_net_(CP; PES) comprises both standard and higher-order components (refer to **section 2.3**). 2) High temporal Pearson correlation (***r***_cp_) observed between CP entropy and the net MI *I*_net_(CP; PES) as well as *S*(PES) and *I*(CP; PES) demonstrates CP-PES phase synchronization. This synchronization is more pronounced under HRG stimulation than under EGF stimulation, as indicated by higher temporal correlations with *S*(CP) (see more details in **Figure 6** and related biological regulations in **section 2.4**). 3) In HRG stimulation, a critical transition occurs within the 10-15-20 min range during the CP-PES phase synchronization, associated with chromatin remodeling and the activation of a Maxwell demon-like mechanism (see **sections 2.5-2.7**). The *x*-axis indicates time points (0, 10, 15, 20, 30, 45, 60, 90, 120, 180, 240, 360, 480, 720, 1440, 2160, 2880, and 4320 minutes), and the *y*-axis represents the measured values (entropy and mutual information in dimensionless units). See the main text for further details.

By replacing *S*(CP,PES) in equation (4) with *S*(WES), the net MI including higher-order terms, *I*_net_(CP; PES) is defined as:

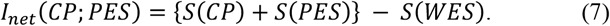

Therefore, from equations (4) and (7), the higher-order MI, *I*_high_(CP; PES) can be defined and observed as positive for the entire experimental time:

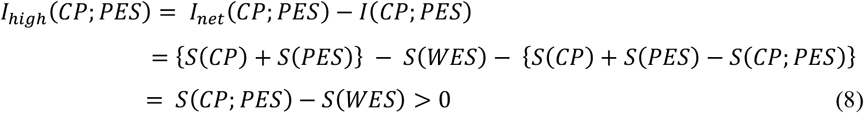

Higher-order MI [McGill, 1954; Hu, 1962; Timme et al., 2014; Rosas et al., 2019] extends beyond standard MI by capturing collective and network-level effects, including overlapping gene functions, feedback loops, and nonlinear interdependencies that simple pairwise correlations cannot fully explain. **In Figure 5**, a positive high-order MI (*I*_high_ >0) under both EGF and HRG stimulation indicates that the total system entropy *S*(WES) is less than the combined CP and PES entropies. This suggests the presence of redundancy (overlapping information) and synergy (collective effects exceeding pairwise contributions) in the MCF-7 genomic network; see partial information decomposition [Williams and Beer, 2010]. These findings support the presence of backup mechanisms, overlapping functions (redundancy), and emergent collective regulation (synergy) in a typical biological genomic network.

In **Figure 6**, under both EGF and HRG stimulation, the net MI, *I*_net_(CP; PES)) are nearly equivalent to the CP entropy S(CP):

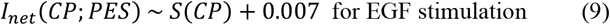

and

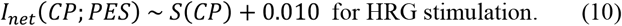

**Figure 6.**
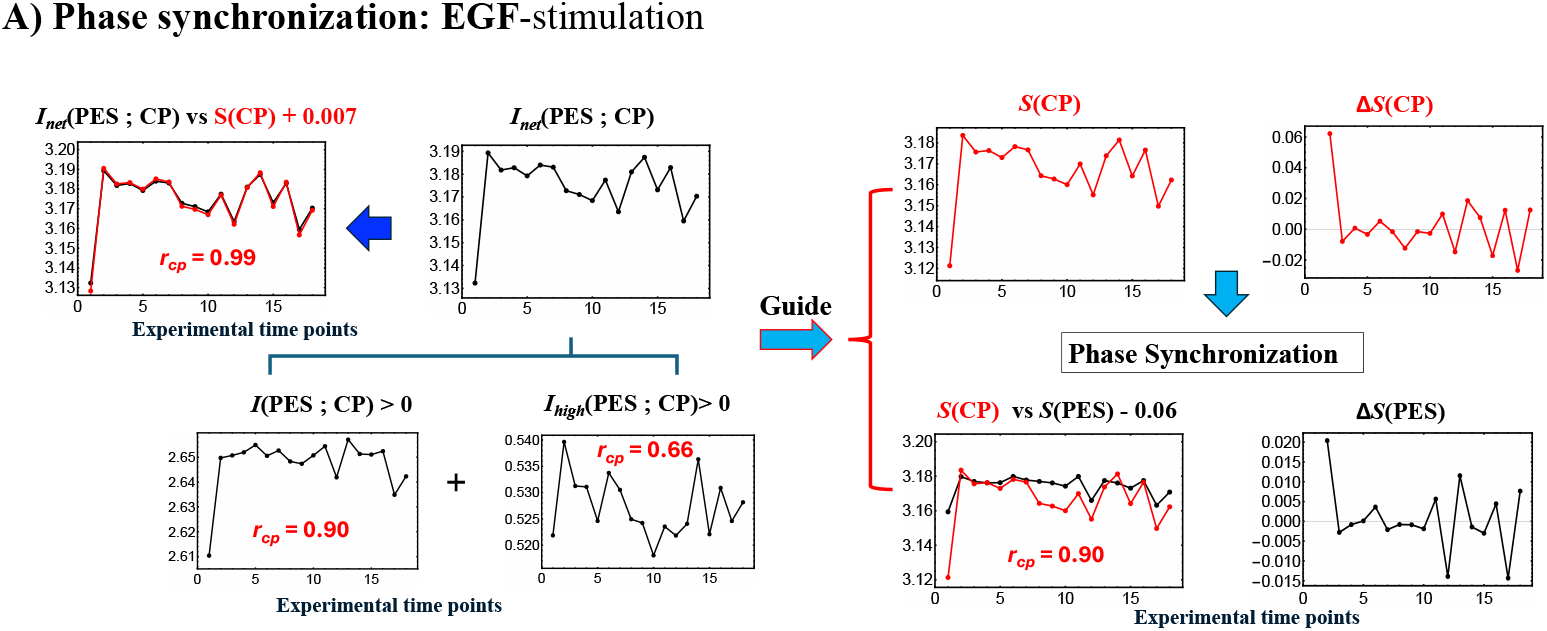
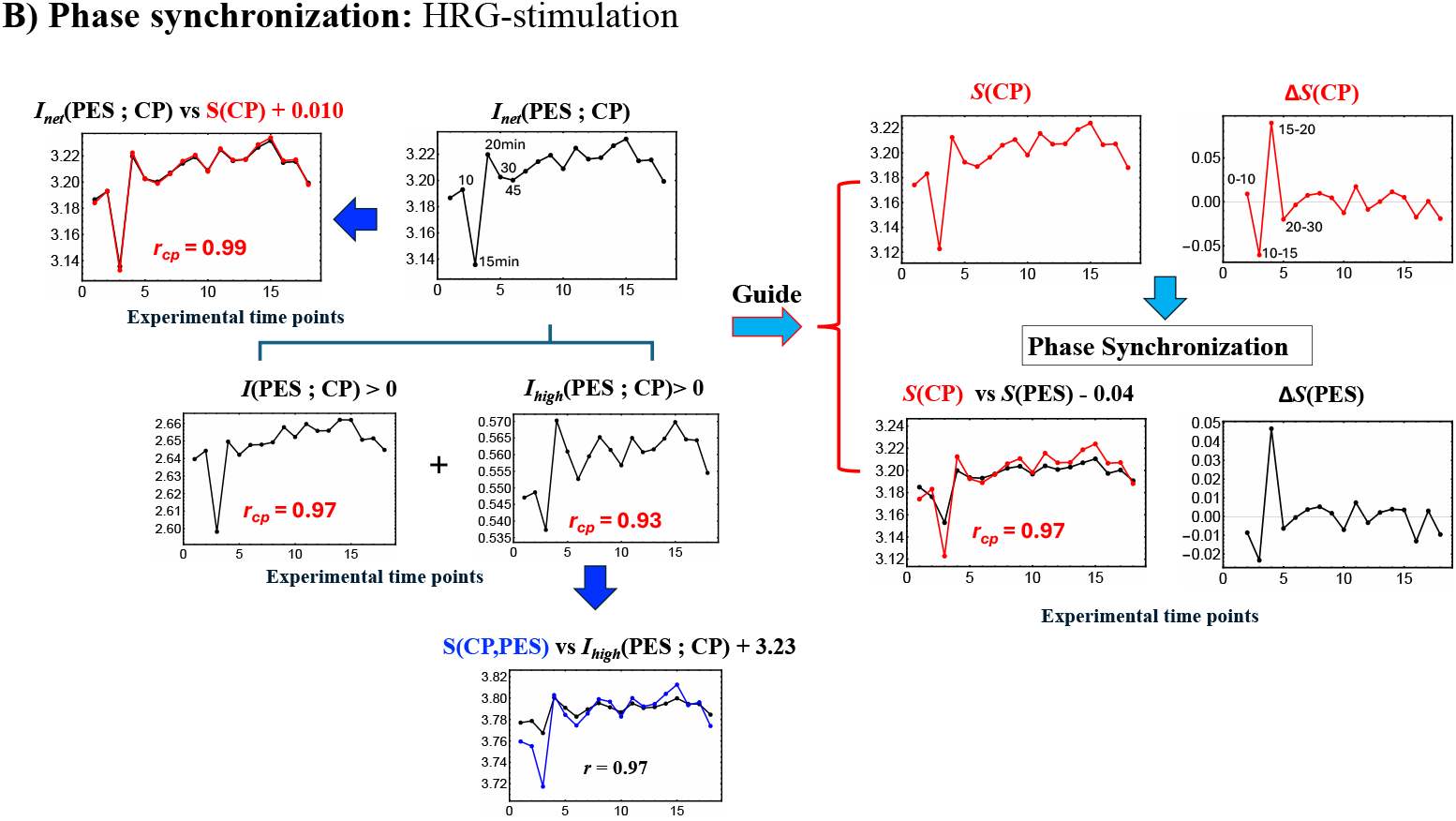
Net MI as Proxy for Thermodynamic CP–PES Phase Synchronization. **A)** EGF stimulation; **B)** HRG stimulation. These figures show that thermodynamic CP-PES phase-synchronization is driven by the net MI, *I*_net_(CP; PES) = *I*(CP; PES) + *I*_high_(CP; PES), where *I*_high_ captures nonlinear contributions from higher-order interactions. In both conditions, *I*_net_ closely follows CP entropy *S*(CP) with a Pearson correlation of ***r***_cp_ = 0.99; under EGF stimulation, *I*_net_ ≈ *S*(CP) + 0.0070, and under HRG stimulation, *I*_net_ ≈ *S*(CP) + 0.010 (upper left panels; black line: *I*_net_; red line: *S*(CP)). For HRG stimulation, *I*_high_(CP; PES), scales with the joint entropy as *S*(CP,PES) ≈ *I*_high_(CP; PES) + 3.23 (***r***_cp_ = 0.97), indicating a substantial nonlinear contribution to CP-PES coupling. During EGF stimulation, the largest entropy changes in S(CP) and S(PES), derived from Inet, occur at 0-10 min, coinciding with early activation of the MAPK/ERK and PI3K/AKT pathways that promote cell proliferation. In contrast, under HRG stimulation, activation of the ERK/Akt pathway at 5-10 min triggers a ligand-specific, biphasic induction of AP-1 components (c-FOS, c-JUN, FRA-1) and c-MYC at 10–20 min, resulting in a pulse-like increase in *I*_net_ from 10 to 30 min. This pulse-like change in *I*_net_ indicates a Maxwell’s demon-like rewritable chromatin memory, as detailed in **sections** 2.5–2.7.

Since the net MI exceeds *S*(CP) slightly, it reflects the positive high-order MI contribution. This result clearly demonstrates that CP genes drive coordinated transitions between ordered and disordered phases between CP and PES. Moreover, it provides a thermodynamic framework to explain how state changes in the CP can drive large-scale ‘genome avalanches,’ as demonstrated through expression flux analysis [Tsuchiya et al., 2023a].

#### 4: Second Law of Thermodynamics

In our open system, the whole expression system comprising CP genes and PES, the entropy production (*σ*) quantifies the irreversible processes occurring within the system. Compliance with the second law of thermodynamics mandates that entropy production must be non-negative (*σ*≥ 0), representing the irreversibility inherent in processes like metabolic reactions, molecular interactions, and active transport mechanisms. Dimensionless entropy production (*σ*) is mathematically defined as the difference between the total change in the WES entropy (Δ*S*(WES)) and the entropy flow exchanged with the external environment of the culture medium (Δ*S*(OUT). Importantly, the net entropy flow, Δ*S*(OUT), is defined with a sign convention where it is positive when the system loses entropy to the environment (i.e., net entropy outflow) and negative when the system gains entropy.

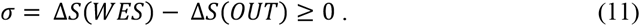

The change in entropy of the external environment, Δ*S*(OUT) is given by

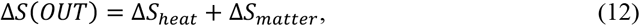

where

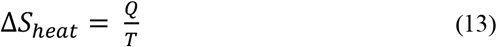

represents the entropy change due to heat (Q) exchange at a temperature (T = 37 °C). By sign convention, Q is negative for heat leaving the system. Δ*S*_matter_ is the net entropy exchange associated with the inflow and outflow of substances:

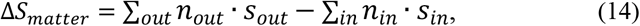

where n_in_ and s_in_ represent the molar quantity and molar entropy of inflowing substances, and n_out_ and s_out_ for outflowing substances.

Cells generate heat through metabolic activities (e.g., ATP synthesis, substrate oxidation), exchange nutrients (e.g., glucose, amino acids) and waste products (e.g., CO_2_, lactate) with the environment.

Therefore, we obtain the second law condition for the system:

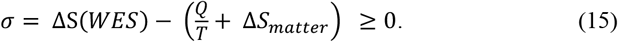

Internal entropy production *σ* must be non-negative (*σ*≥ 0) to satisfy the second law of thermodynamics. This ensures that all irreversible processes within the cell culture contribute to an overall increase in entropy.

### 2.4 Genomic-Thermodynamic Phase Synchronization Between CP Genes and the Whole Expression System

The genomic-thermodynamic phase synchronization dynamics for EGF-stimulated and HRG-stimulated MCF-7 cells is summarized below:

#### EGF-stimulated MCF-7 cells (no cell fate change)

EGFR activation drives entropy changes in CP and PES genes within the first 10 min. Afterward, information-thermodynamic metrics fluctuate around average values through synchronized order-disorder phases, guided by net mutual information dynamics scaling with CP entropy. WES maintains a dynamic balance of entropy and information flux.

#### HRG-stimulated MCF-7 cells (cell fate change)

PES exhibits stronger phase synchronization with CP than under EGF stimulation via net MI dynamics. A critical transition at 10-30 min activates the CP as a Maxwell demon, driving feedback to the PES and triggering genomic avalanche. This leads to a pulse-like perturbation in all information-thermodynamic metrics.

A key difference between EGF and HRG stimulation is the emergence of a critical transition under HRG, driven by the activation of a Maxwell’s demon-like mechanism (**sections 2.5** and **2.6**; see also the conclusive remarks in **section 2.7**).

Further details are provided below:

##### 1) In EGF-stimulated MCF-7 cells

###### A) Dissipative Thermodynamic Synchronization of Order-Disorder Phases

In EGF-stimulated MCF-7 cells, EGFR activation triggers the MAPK/ERK pathway for gene regulation and the PI3K/AKT pathway for survival, driving proliferation within the first 10 min [Saeki et al., 2009]. This activation induces marked changes in the entropy of the CP region and the PES, as evidenced by the rise in net MI, *I*_net_(CP;PES), during the initial 0-10 min (**Figure 5A**).

The increase in *I*_net_(CP;PES) reflects nonlinear interactions between CP and PES that expand both of their accessible state microspaces, which indicates active exchanges of heat and matter with the environment. As the CP’s complexity grows, its nonlinear influence on the PES strengthens, promoting greater integration between these subsystems. Furthermore, the phase synchronization between CP and PES, characterized by coordinated shifts between ordered and disordered phases (**Figure 6A**), yields a high Pearson correlation (r = 0.90). This synchronization is fundamentally driven by *I*_net_(CP;PES), which combines standard MI with positive higher-order MI (**section 2.3**) and scales with S(CP).

###### B) Dynamic Equilibrium Underpinning Coherent Stochastic Behavior (CSB) In Figure 5A

*S*(WES) is notably stable, ranging from approximately 3.208 to 3.234, with a total variation of about 0.026 units around its mean of 3.227. This balance of entropy and information flux preserves overall stability and supports coherent stochastic behavior (CSB). This observation aligns with expression flux analyses [Tsuchiya et al., 2020, 2023a], suggesting a biophysical correlate of the law of large numbers, where the collective system behavior remains stable despite the inherent stochasticity of individual molecular interactions.

##### 2) In HRG-stimulated MCF-7 cells

###### Activation of the CP as a Maxwell Demon

In contrast to the large initial disorder changes induced by EGF in the first 0–10 min, HRG stimulation triggers pulse-like order-disorder transitions in all information-thermodynamic metrics, including both S(CP) and S(PES) as net MI *I*_net_(CP;PES) rises during the 10–20 minute period (**Figure 5B**). This suggests more active exchanges of heat and matter with the environment under HRG, strengthening phase synchronization. The pulse-like critical transition aligns with findings from PCA [Zimatore et al., 2021] and expression flux analysis [Tsuchiya et al., 2020, 2022]. It involves the CP acting as a Maxwell demon (see detailed analysis in later sections), inducing feedback to the PES and orchestrating a genomic avalanche that drives cell-fate change.

From a biological standpoint, **within 5-10 min t**ime window, HRG stimulation, distinct from EGF, activates the ERK/Akt pathway, leading to a ligand-specific, biphasic induction of AP-1 complex components (e.g., c-FOS, c-JUN, FRA-1) and the transcription factor c-MYC [Nagashima et al., 2007; Saeki et al., 2009; Nakakuki et al., 2010].

###### By 15-20 min

the above-sketched biochemical cascade produces a pulse-like genome-wide perturbation, marked by the activation of oscillating-mode genes (OMGs), as shown by expression flux analysis [Tsuchiya et al., 2023a]. These genes emit large expression fluxes to both the CP and the genome attractor (GA), thereby facilitating their synchronization within the OMG–CP–GA network.

###### At 60 min

this critical transition is followed by a peak in c-MYC expression [Krigerts et al., 2021; Erenpreisa et al., 2023], resulting in extensive gene amplification. Beyond this point, entropy changes exhibit steady fluctuations with damped behavior, as observed also in both PCA and expression flux analyses. This indicates balanced large-scale energy flows that sustain cellular homeostasis.

These findings on EGF- and HRG-stimulation demonstrate that changes in entropy and net mutual information effectively serve as indicators of fundamental cellular processes, including the activation of signaling pathways, the occurrence of critical transitions, and the synchronization of genome-wide expression. Adopting this thermodynamic perspective enriches our understanding of how different stimuli orchestrate complex regulatory networks within cells.

### 2.5 CP Acts as Maxwell’s Demon in Early HRG Signaling Response

Unlike EGF, which induces early disorder (0-10 min), HRG triggers a distinct critical transition, synchronizing ordered and disordered phases across all information-thermodynamic metrics from 10 to 20 min. As shown in **Figures 5B** and **6B**, the CP acts as a Maxwell’s demon [Parrondo et al., 2015], regulating information flow and maintaining adaptable chromatin states (see **section 2.6** for details).

The CP, functioning as a Maxwell’s demon, orchestrates gene expression and cell fate through the following three distinct phases:

#### Phase 1: Preparation (10-15 min)

The CP synchronizes with the PES to minimize the entropies of both CP and PES genes, establishing their initial states. Biologically, this synchronization sustains the upregulation of AP-1 complex genes and coordinates the activation of their downstream targets, ensuring proper assembly and functionality of AP-1 components (see **section 2.4**).

#### Phase 2: Measurement (15-20 min)

Synchronization between the CP and PES initiates the measurement process, leading to the largest increases across all information-thermodynamic metrics, including CP and PES entropy, as well as standard, higher-order, and net MI.

#### Phase 3: Feedback and Critical Transition (20-30 min)

Utilizing the net MI obtained in Phase 2, the CP reorganizes and provides feedback to the PES, triggering a critical transition around 20 min in the WES [Tsuchiya et al., 2020, 2022; Zimatore et al., 2021]. This feedback-driven reorganization reduces system entropy, driving the system into a new ordered state.

Further details are provided below:

### Phase 2: Measurement Phase (15–20 min)

#### 1. Highest Increases in CP Entropy, PES Entropy, and Net MI

As shown by the 15- and 20-minute data points in **Figure 5B** (or **Figure 6B**), CP genes acquire the most information about the PES during the measurement phase. Processing and storing this information raises CP entropy due to thermodynamic costs. This phase sees the largest increases in both standard and higher-order MI, driving net MI from its 15-minute minimum. Notably, the surge in higher-order MI underscores strong nonlinear interdependencies between CP and PES.

**Figure 6B** shows that higher-order MI strongly correlates with the joint entropy *S*(CP,PES) (*r* = 0.97). As total uncertainty increases, more states become accessible, broadening the system’s global search space and amplifying structured interactions between the CP and PES. The positive higher-order MI reflects both synergy (emergent collective information) and redundancy (overlapping information) (see **section 2.3**), enabling the CP to encode information about the PES despite rising global uncertainty. This open thermodynamic phase synchronization, led by CP genes, is a hallmark of complex systems, where localized organization persists in high-entropy environments, supporting robust information processing.

**Note:** In the scenario described by Parrondo et al. (2015), the entropy of the measured system (here, the PES) decreases as uncertainty is transferred to the measuring agent (the CP), thereby increasing the system’s internal order. In contrast, in our open CP-PES synchronized system, the PES, along with the CP, continuously exchange energy with the environment. During phase synchronization, instead of experiencing a net entropy reduction due to information transfer, the PES absorbs energy, such as heat, free energy, or chemical substrates, from its surroundings. This energy absorption offsets the measurement-induced entropy decrease and produces a net increase in *S*(PES). Consequently, this continuous energy influx sustains phase synchronization between the CP and PES and supports the feedback mechanism crucial for Maxwell’s demon function.

The synchronized metrics in Phase 2 lay the foundation for the feedback process in Phase 3, driving the system’s critical transition to a new cell state.

### Phase 3: Feedback and Critical transition

#### 1. Initiation of Critical Transition (20-30 min)

During this interval, the CP’s decreasing entropy induces a corresponding drop in PES entropy through phase synchronization, facilitated by exchange entropy and information flow with the environment. Through feedback, the CP utilizes net MI to drive a critical transition, revealing the thermodynamic costs of its activity. Concurrently, declines in both standard MI and higher-order MI reduce net MI, suggesting that the CP is leveraging information from Phase 2 to guide the PES through a critical transition. Notably, the timing of this CP feedback around 20 min aligns with a genome-wide transition, as shown by expression flux analyses [Tsuchiya et al., 2023a] and PCA [Zimatore et al., 2021].

#### 2. Stabilization into New States After 30 min

After 30 min, fluctuations in *S*(CP), *S*(PES), and net MI suggest that both the CP and PES are settling into stable, reorganized states. Notably, between 30 and 45 min, the bimodal CP dissipates, and by 60-90 min, a new CP emerges, indicating the establishment of a new state (see **section 2.6**).

### 2.6 Maxwell’s Demon Functioning as Rewritable Chromatin Memory

J. Krigerts et al. (2021) experimentally investigated critical transitions in chromatin dynamics during early differentiation in HRG-stimulated MCF-7 cells, focusing on pericentromere-associated domains (PADs). They identified two key critical transitions:

#### 1. First Critical Transition (15-20 min)

Following HRG treatment, PADs undergo a “burst,” dispersing from their clustered state near the nucleolus. This dispersal coincides with the activation of early response genes, such as c-fos, fosL1, and c-myc, which initiate the differentiation process. During this phase, repressive chromatin structures unravel, and active chromatin regions become more accessible, marking a significant step in genome reorganization (see **section 2.7** for power-law scaling of PAD Distribution).

#### 2. Second Critical Transition (Around 60 min)

The second transition involves further chromatin alterations, including increased transcription of long non-coding RNAs (lncRNAs) from PADs. This supports additional chromatin unfolding and genome rewiring, which are essential for establishing and maintaining differentiation, ultimately determining a stable and functional cell fate.

In this section, we explore the information-thermodynamic mechanisms underlying these two critical transitions in chromatin dynamics. To achieve this, we analyze the metric *ln*(*nrmsf*) (equation (1)), which indicates the self-organization of the whole expression system (WES) into distinct response domains separated by a critical point (CP) (see **Figure 2**; refer to [Tsuchiya et al., 2023b] for details). This metric quantifies the temporal variability of gene expression as a time-independent measure and serves as a proxy for chromatin remodeling dynamics.

Zimatore et al. (2021) showed that *ln*(*nrmsf*) accurately captures the temporal variability of gene expression and that principal component (PC) scores derived from whole transcriptome time series data predict each gene’s *ln*(*nrmsf*) (see **section 2.1**). During the critical transition between 10 and 30 min, the variance explained by the second principal component (PC2) increases by one order of magnitude among oscillating mode genes (OMGs) [Tsuchiya et al., 2023a], delineating the primary axis of deviation from the genome attractor (GA). This amplification of PC2 coincides with pericentromeric body splitting and chromatin unfolding, events that expose previously inaccessible genomic regions to transcriptional machinery [Krigerts et al., 2021] (see **section 2.7**). Because PC2 dynamics mirror *ln*(*nrmsf*) values and Shannon entropy is computed directly from the *ln*(*nrmsf*) distribution, *ln*(*nrmsf*) serves as a quantitative proxy for entropy changes, thereby mechanistically linking chromatin remodeling to thermodynamic shifts.

As illustrated in **Figure 7**, raw gene expression data are binned into *m* discrete *ln*(*nrmsf*) intervals that define “logical states”, corresponding to chromatin states representing broad, coarse-grained chromatin conformations (e.g., transitions between more compact and more open structures), while the expression values within each interval define “physical states”, capturing finer scale variability in gene expression within each logical state.

**Figure 7.**
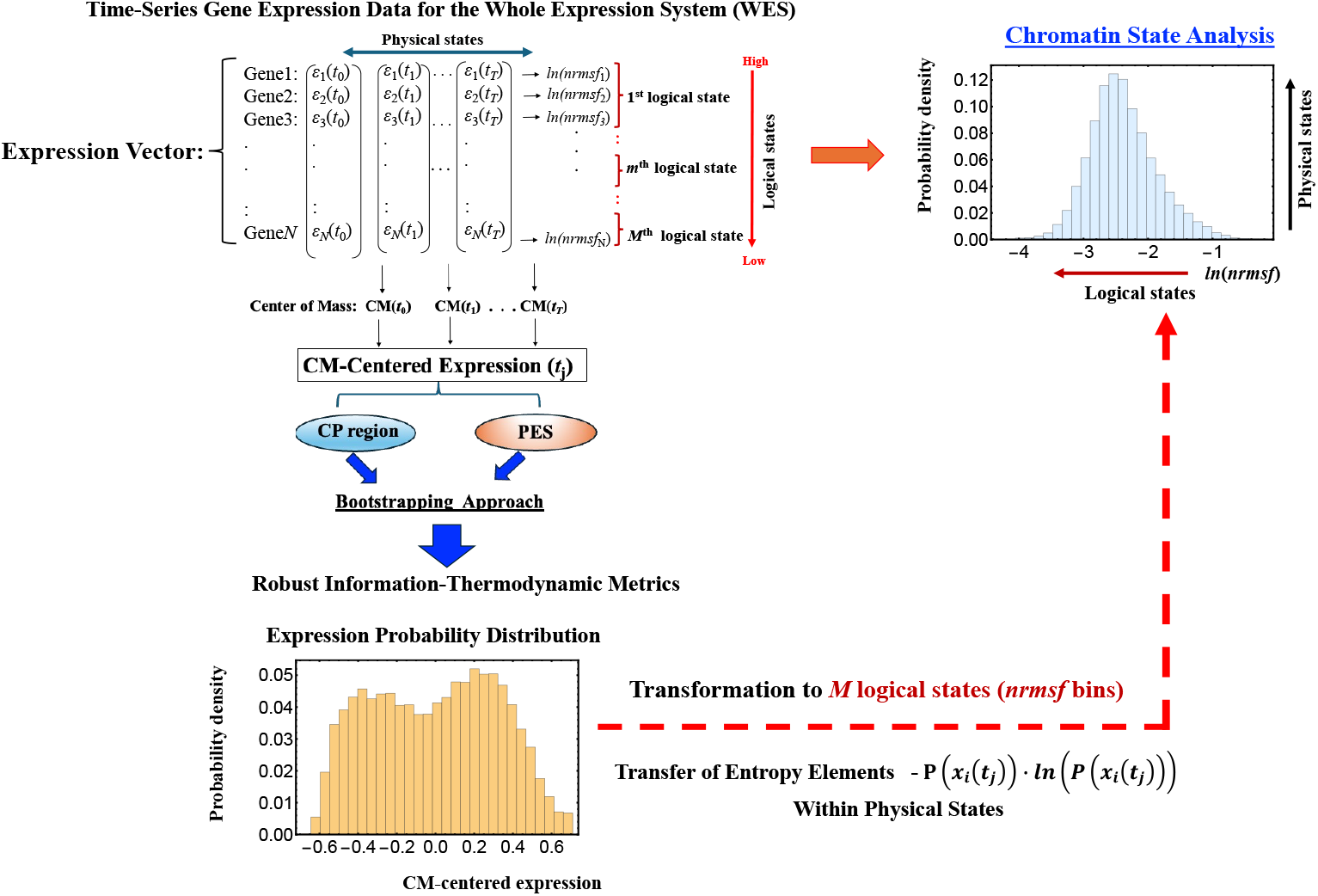
Information-Theoretic Workflow for Converting Gene Expression to *ln*(*nrmsf*) Probability Distributions. This schematic outlines the procedure for transforming gene expression probability distributions into *ln*(*nrmsf*) probability distributions. In the *ln*(*nrmsf*) probability distribution, the *x*-axis represents logical states, defined as *ln*(*nrmsf*) (fixed) bins that quantify large-scale transitions in chromatin states (see **sections 2.6** and **2.7**), while the *y*-axis captures physical states, reflecting the fine-scale variability of gene expression within each chromatin domain. The workflow begins with temporal gene expression data from the WES, which encompasses both the CP region and the PES. This approach ensures the robustness of our information-thermodynamic metrics (see **Figure 1**). Entropy components, given by - *p*(*x*_i_(*t*_j_))*ln*(*p*(*x*_i_(*t*_j_))), are computed at each time point *t*_*j*_ and then mapped onto the *ln*(*nrmsf*) bins, thereby converting detailed gene expression profiles into logical states that highlight major chromatin state transitions. This transformation allows us to capture dynamic variability at both the logical (large-scale) and physical (local) levels. For additional details on this process, refer to **Figure 8**.

This hierarchical structure reflects the entropy decomposition into logical and physical components, as proposed by Maroney (2009) in his extension of Landauer’s principle. In this framework, the entropy of the logical states quantifies large-scale chromatin reorganization, while the residual entropy of the physical states captures gene-specific regulatory fluctuations embedded within each chromatin domain.

Robust entropy estimates are obtained with the convergence protocol detailed in **Figures 1 and 4**. The elemental term -*p*(*x*_i_(*t*_j_))*ln*(*p*(*x*_i_(*t*_j_))) is iteratively assigned to its corresponding *ln*(*nrmsf*) bin, and bootstrapping is repeated until the convergence criterion is satisfied. This procedure ensures that each expression value is fully incorporated at the logical level and that the resulting entropy estimates are both numerically stable and reproducible.

By integrating logical and physical components, our ITA links abrupt structural transitions, such as the sharp increase in variance captured by PC2, to their effects on gene-expression landscapes. The analysis therefore clarifies how chromatin dynamics modulate transcriptional states and ultimately steer cell-fate decisions.

Note that the workflow in **Figure 7** analyzes chromatin dynamics using a fixed number of chromatin states (*ln*(*nrmsf*) bins). The two-dimensional approach in the **Appendix** extends this framework by simultaneously incorporating logical and physical states, enabling an integrated assessment of both large-scale and local entropy contributions.

**Figure 8** depicts the temporal variations in Shannon entropy linked to chromatin dynamics across the WES for EGF- and HRG-stimulated MCF-7 cells:

**Figure 8:**
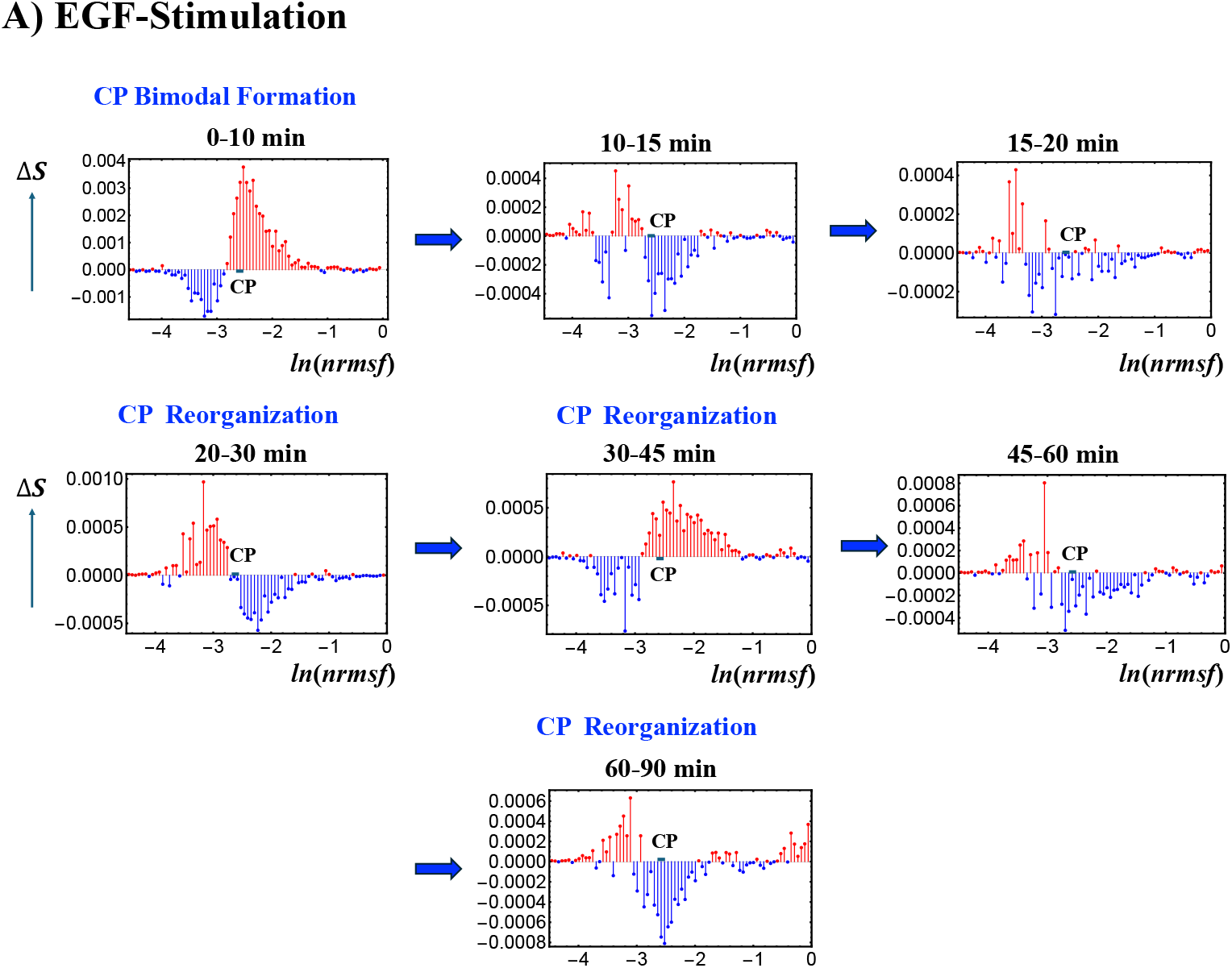
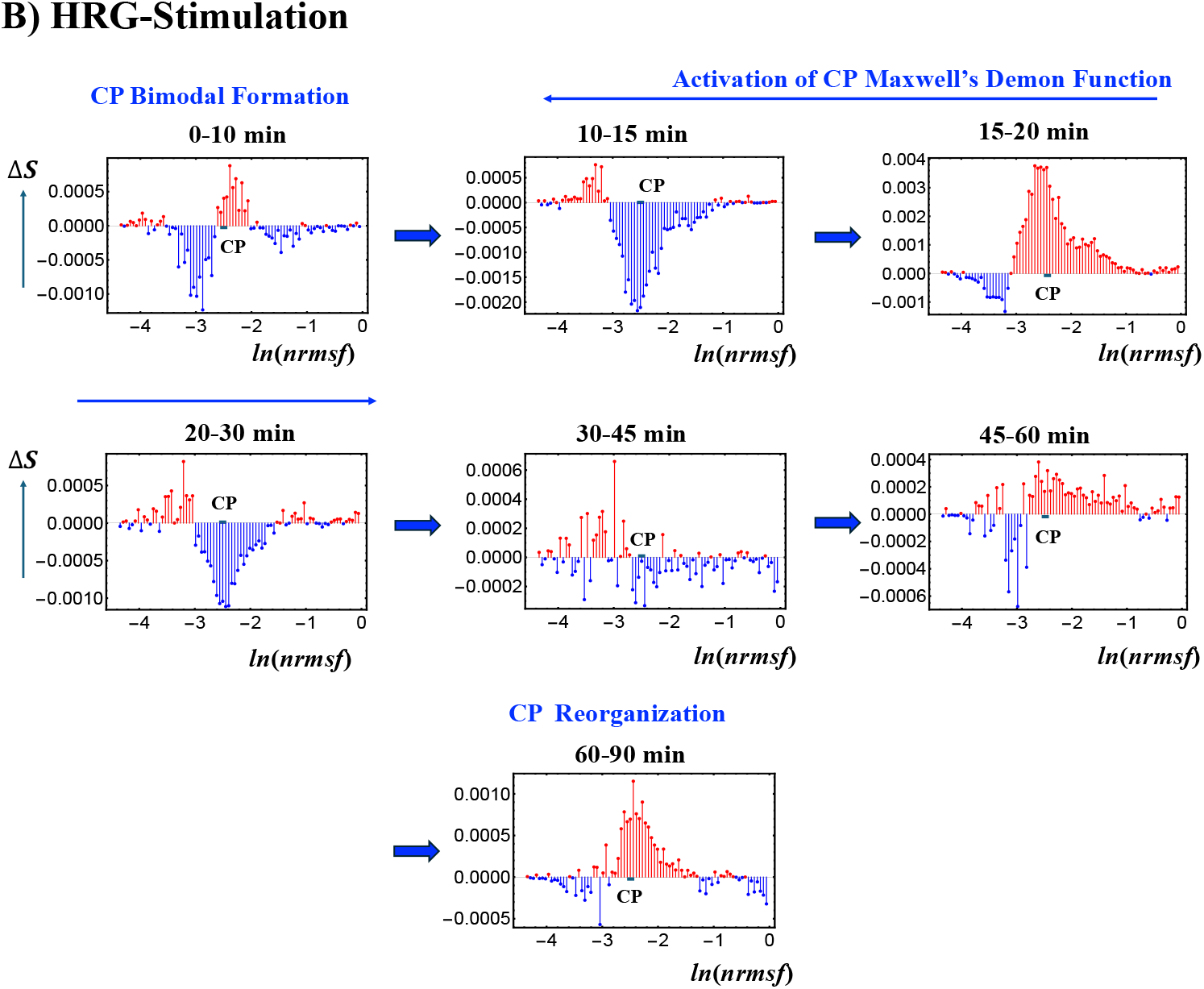
Chromatin Dynamics and CP Bimodal Formation Across Temporal Entropy Changes. Temporal changes in Shannon entropy, reflecting chromatin dynamics, are plotted against the natural logarithm of *nrmsf* (in 80 bins) **A)** in EGF- and **B)** HRG-stimulated MCF-7 cells. The CP genes, marked by the bold black solid line, displays singular bimodal behavior within the ranges -2.64 < *ln*(*nrmsf*) < - 2.52 for EGF stimulation and -2.55 < *ln(nrmsf*) < -2.44 for HRG stimulation (see **Figure 2**). Under EGF stimulation, intermittent CP bimodal patterns emerge at specific intervals (e.g., 0-10 and 20-30 min) but repeatedly dissolve without triggering a critical transition, indicating that coherent global chromatin remodeling does not occur, resulting in no cell-fate change.

#### 1. EGF-stimulated MCF-7 cells (Figure 8A)

This panel shows the temporal changes in Shannon entropy linked to chromatin dynamics in EGF-stimulated MCF-7 cells. Within the specified range (−2.64 < *ln*(*nrmsf*) < -2.52; **Figure 1**), a CP-like (critical point-like) bimodal pattern in entropy changes occasionally appears, suggesting transient critical behaviors in chromatin organization. During these intervals, positive entropy changes (ΔS > 0) are associated with chromatin unfolding, while negative entropy changes (ΔS < 0) are associated with chromatin folding.

Although these CP-like bimodal patterns emerge intermittently, such as during the 0-10 and 20-30 min windows, they dissolve and reappear without inducing a sustained critical transition or altering cell fate. This implies that the conditions needed to activate a Maxwell’s demon-like mechanism are not fully met. Notably, these chromatin fluctuations are driven by CP–PES phase synchronization, underscoring the role of thermodynamic interactions in maintaining dynamic chromatin states despite the absence of a full critical transition.

#### 2. HRG-stimulated MCF-7 cells (Figure 8B)

The scenario of Maxwell’s demon-like chromatin memory progresses through distinct phases. During the initial 0-10 min, a CP-like bimodal pattern (−2.55 < *ln*(*nrmsf*) < -2.44; **Figure 1**) emerges. At 10-30 min, a global fold-unfold coherent transition occurs around *ln*(*nrmsf*) = -3.11 to -3.04, involving Maxwell’s demon-like behavior (see **section 2.5**). This phase acts as a rewritable chromatin memory, reflecting changes in double-well potential profiles (**Figure 9**). At 30-45 min, the CP pattern dissolves, likely preparing the system for a new stable genomic state of the WES. At 60-90 min, CP formation reappears, suggesting a cell-fate change consistent with the second transition described by Krigerts et al. (2021). Beyond 90 min, CP-like bimodal patterns continue to form and dissolve intermittently, indicating sustained chromatin adaptation under HRG stimulation.

**Figure 9:**
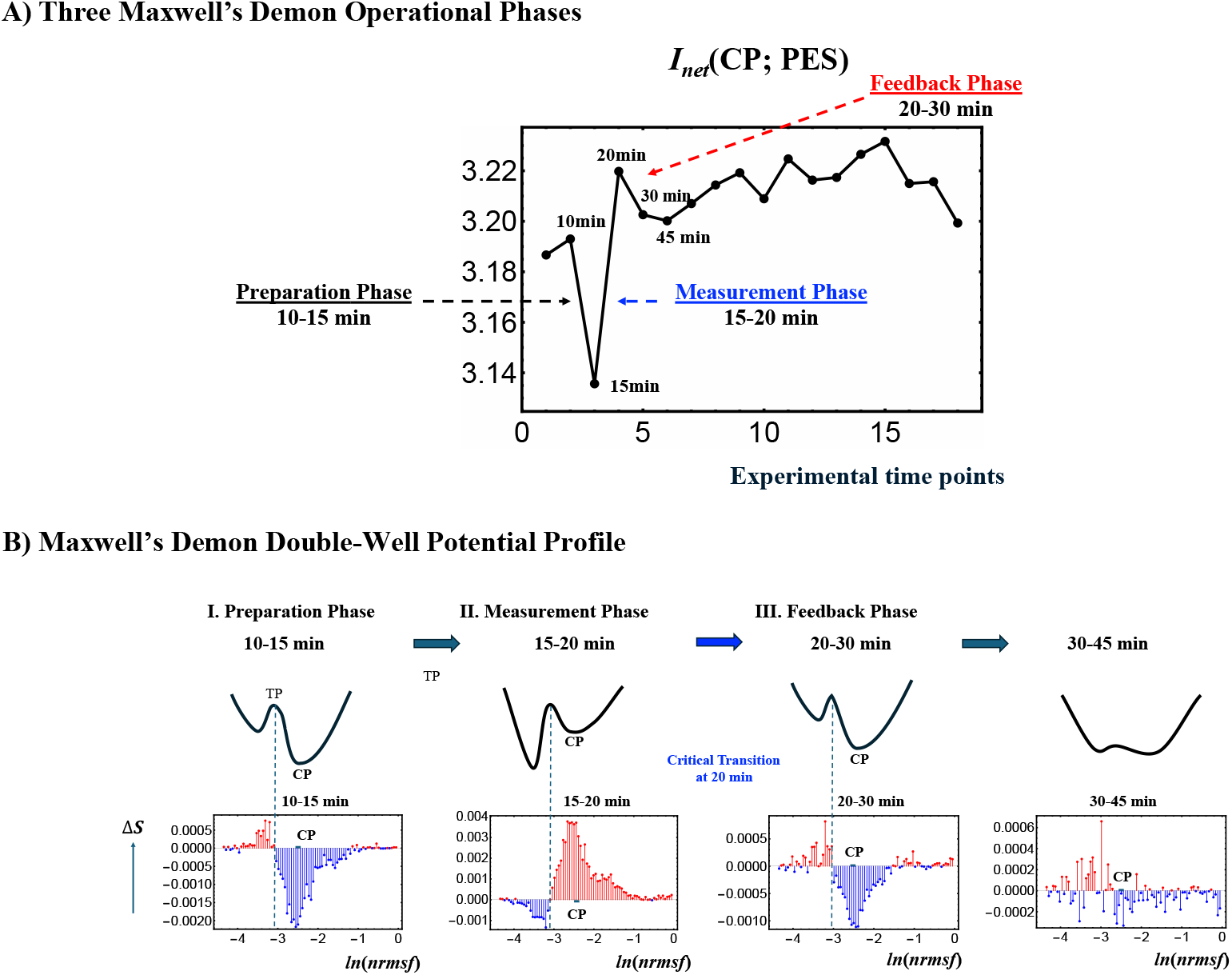
Three Maxwell’s Demon Processes and Schematic Representation of Its Double-Well Potential Profile. **A)** Net MI *I*_net_ (CP;PES) in HRG stimulation reveals all three operational phases of a Maxwell’s demon: preparation (10-15 min), measurement (15-20 min), and feedback (20-30 min). **B)** This panel illustrates a conceptual double-well potential landscape representing chromatin dynamics, highlighting the Maxwell’s demon-like behavior of the CP within the range -2.55 < *ln*(*nrmsf*) < -2.44. Each well corresponds to a distinct chromatin state, such as folded or unfolded chromatin, separated by a potential barrier. The transition point (TP) occurs around *ln*(*nrmsf*) = -3.15 to -3.00, marking where chromatin restructuring takes place. Chromatin remodeling dynamics are visually represented by up (red) and down (blue) arrows, reflecting the oscillatory behavior of coherent chromatin folding and unfolding, dynamically centered around the CP. After 30 min, the potential barrier almost disappears, suggesting an inactive Maxwell’s demon function with a loss of coherent chromatin state. **A)** The *x*-axis represents experimental time points, and the *y*-axis represents the net MI value. B) The *x*-axis represents changes in Shannon entropy associated with chromatin remodeling, and the *y*-axis represents the value of *ln(nrmsf*).

Using the *ln*(*nrmsf*) metric, Shannon entropy reveals chromatin folding and unfolding dynamics. EGF stimulation repeatedly dissolves and reinitiates CP bimodal pattern formations without inducing cell fate changes, while HRG stimulation facilitates stable CP formation and cell fate transitions driven by Maxwell’s demon-like chromatin memory at 10-30 min. The CP’s role, measuring genomic states, orchestrating entropic and informational dynamics, and reorganizing chromatin, demonstrates its active function in encoding, decoding, and re-encoding chromatin configurations. This process aligns with the concept of memory storage and rewriting within a dynamically regulated, thermodynamically controlled genomic landscape (**Figure 9**).

In contrast, HRG stimulation initially produces a CP bimodal pattern during the 0-10 min period, which then transitions into a global, coherent fold-unfold chromatin change between 10-30 min (see **section 2.7** for experimental evidence). This is followed by the re-formation of CP states between 60-90 min, aligning with a cell-fate transition (see **section 2.6**). These results highlight the emergence of cyclical chromatin organization states, driven by CP-PES phase synchronization and information-thermodynamic interactions, including Maxwell’s demon-like memory effects in chromatin structure (see more detail in **Figure 9**).

### 2.7 Structural Signatures of Chromatin Remodeling in HRG-stimulated MCF-7 Cells

Here, we provide further experimental evidence supporting the use of gene expression variability, quantified by *ln*(*nrmsf*) values, as a proxy for chromatin remodeling dynamics.

Chromatin folding and unfolding are highly complex biochemical processes, underscored by the enormous challenge of compressing approximately 2 meters of human DNA into the few-micrometer space of a cell nucleus. The “structural signatures” of chromatin remodeling were revealed through the combined application of confocal microscopy image analysis and biochemical approaches [Krigerts et al., 2021]. In their study, Krigerts and colleagues focused on Pericentric-Associated Domains (PADs) and chromocentres, higher-order complexes formed by the aggregation of PADs.

Chromocentres serve as markers of chromatin’s folded or unfolded state, co-segregating with repressive genomic regions and contributing to gene silencing near centromeres via position effect variegation [Bártová et al., 2002; Probst and Almouzni, 2008; Carelli et al., 2017]. PADs, and consequently chromocentres, form through the transient aggregation of histones and protamines, and are characterized by a highly variable acetylation pattern [Probst and Almouzni, 2008]. The relative size of chromocentres determines their effect on chromatin density. A single PAD has an approximate area of 1 µm^2^. Chromocentres composed of fewer PADs indicate lower chromatin density, which is associated with increased gene expression variability.

In the early phase of the cell fate transition in HRG-stimulated cells, corresponding to 15-30 min after drug administration, a marked disaggregation of chromocentres is observed, giving rise to isolated PADs. This structural transition aligns with the critical transition described in **sections 2.4-2.6**. Furthermore, as expected in an aggregation-disaggregation process, the number of chromocentres follows a power-law scaling with their relative size (see **Figure 10**).

**Figure 10.**
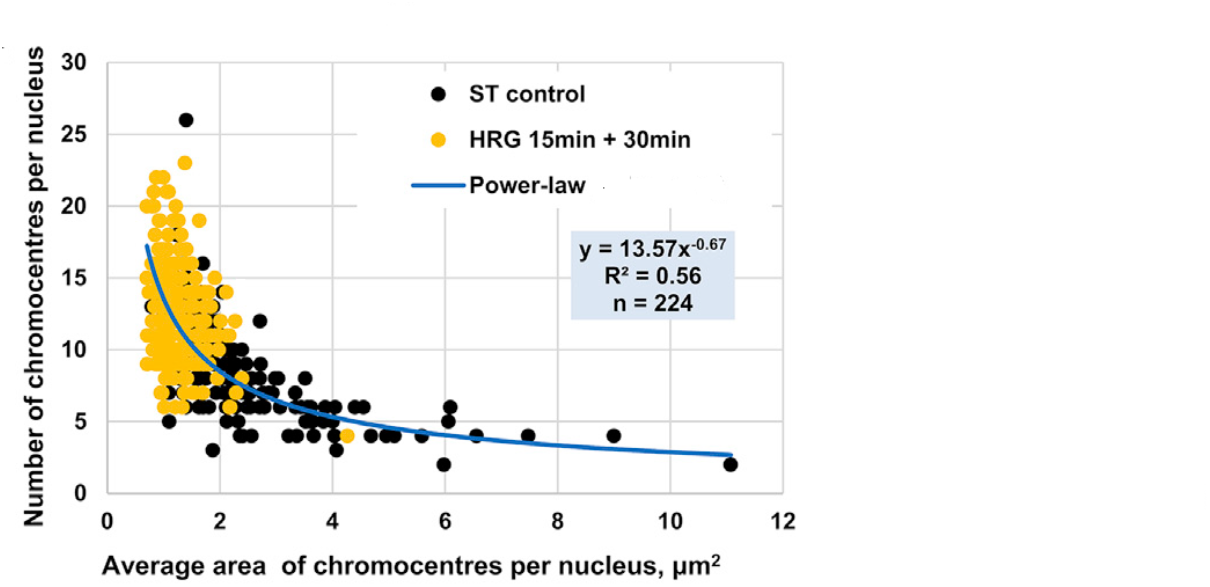
Power-Law Scaling of Chromocentre Distribution in MCF-7 Cells Under HRG Stimulation. Area (*x*-axis) and number of PADs (*y*-axis) in MCF-7 cells as observed by confocal microscopy. Black dots represent PADs in untreated (ST control) cells, while yellow dots indicate PADs under HRG stimulation during the transition phase. The distribution follows a power-law scaling: small chromocentres significantly outnumber larger ones, with most chromocentres comprising a single PAD during the critical transition.

During the transition phase, chromocentres disaggregate into single PADs, corresponding to targeted chromatin unfolding and a consequent increase in gene expression variability. The dynamics of PADs provide proof-of-concept for the hypothesis that temporal gene expression variability by *ln*(*nrmsf*) values is a functional consequence of chromatin remodeling. This notion is further supported by the observed upregulation of key differentiation master genes such as FOS and c-Myc [Krigerts et al., 2021].

Together, these findings establish a crucial link between our statistical mechanics framework, which views the genome as a unified system whose global dynamics are reflected in the distribution of gene expression variability, and the classical, single-gene perspective of molecular biology (see also **Discussion**).

As a conclusive remark, our information-thermodynamics analysis (ITA) demonstrates that the CP functions as a Maxwell’s demon in genome regulation by driving cell-fate changes. This conclusion is supported by the following four independent lines of evidence:

1. The CP fulfills all three operational phases characteristic of a Maxwell’s demon [Sagawa and Ueda, 2012; Parrondo et al., 2015] (**section 2.5**).
2. Experimental and computational analyses of chromatin dynamics based on *ln*(*nrmsf*)-sorted whole-genome expression data consistently reveal regulatory patterns indicative of Maxwell’s demon–like behavior (**sections 2.6** and **2.7**).
3. These findings are confirmed by an independent replicate of the gene expression dataset of HRG-stimulated MCF-7 cells (see **Material**) as detailed in the **Supplemental File**.
4. Similar regulatory behavior observed in a different cancer cell type, DMSO (dimethyl sulfoxide)-stimulated HL-60 cells [Huang et al., 2005], as detailed in the **Supplemental File**, further reinforces this conclusion.

## 3. Discussion: Biophysical Implications

Our findings highlight the genome as an open thermodynamic system operating far from equilibrium, where continuous energy dissipation sustains dynamic chromatin organization and gene expression. This dissipative framework explains how cells preserve functional stability amid environmental fluctuations.

From a biomedical standpoint, this thermodynamic perspective has substantial implications. By elucidating the principles governing how cells adopt and maintain distinct fates, we gain insights into processes such as cancer progression, metastasis, and therapeutic resistance. Targeting the CP hub may therapeutically reprogram cell states by altering chromatin remodeling, gene expression, signaling, epigenetics, cell cycle and metabolism. For instance, restricting undesirable plasticity in cancer cells or guiding stem cells toward more favorable differentiation pathways could offer new strategies for improved therapeutic interventions.

In Discussion section, we further expand our findings and discuss the potential for controlling the dynamics on the fate of cancer cells in three key aspects: 1) order parameter of phase synchronization, 2) autonomous genome computing and 3) genome intelligence (GI).

### 1) Existence of an Order Parameter for Phase Synchronization by Integrating Dynamical System Analysis and Non-Equilibrium Thermodynamics

We explore the emergence of an order parameter for phase synchronization by integrating the genome engine mechanism, a dynamical systems approach based on expression flux analysis (summarized in the Introduction and detailed in [Tsuchiya et al., 2020, 2022, 2023a]), with genomic thermodynamic phase synchronization, developed as a non-equilibrium thermodynamic framework in this study. This integration suggests a biophysical link between dynamical principles and genome regulatory mechanisms, leading to the emergence of a measurable order parameter for phase synchronization.

Under HRG stimulation, as illustrated in **Figure 5B**, the net mutual information between the CP and the PES, reflecting nonlinear higher-order interactions, drives the emergence of CP-PES thermodynamic phase synchronization. In particular, the critical transition triggers a drastic system-wide shift, where all statistical measures describing the state become highly correlated, effectively collapsing the dynamical space into a binary regime: CP activation under HRG versus no CP activation under EGF. As a result, every computed index converges within a unified critical framework. This indicates that the dynamics of phase transitions not only drive biological coherence, as exemplified by the pulse phase in biphasic induction of the AP-1 complex [Nakakuki et al., 2010], but also integrate diverse analytical metrics into a single, self-organizing system.

**Figure 10** provides direct evidence of this collapse, showing that the CP entropy change (ΔS(CP)), representing thermodynamic behavior, synchronizes with the temporal evolution of self-flux (effective force) at both the CP and the genome attractor (GA), thereby guiding large-scale changes in genomic expression dynamics (see **Introduction**). This concurrent synchronization bridges thermodynamic and dynamical system behaviors. Formerly independent statistical descriptors become tightly correlated, and simultaneous surges in ΔS(CP), CP self-flux, and GA self-flux confirm that thermodynamic and dynamical cues act synergistically to enforce genome-wide phase coherence.

The convergence of different descriptors suggests the emergence of a robust global order parameter, analogous to the Kuramoto measure [Kuramoto, 1975; Strogatz, 2000]. This scalar quantity, ranging from near zero (indicating incoherent or random phase relationships) to near one (denoting near-perfect synchrony), encapsulates the degree of phase alignment between the CP and GA. As the CP and GA become increasingly synchronized, the order parameter rises steeply, marking the transition from a disordered to an ordered state, a signature hallmark of self-organized critical (SOC) control in the whole expression system. Thus, this pulse-like phase transition not only underscores the CP’s role as the central regulatory hub in cancer cell expression but also bridges dynamical and statistical perspectives, unifying diverse analytical metrics into a single, self-operating system.

**Figure 10.**
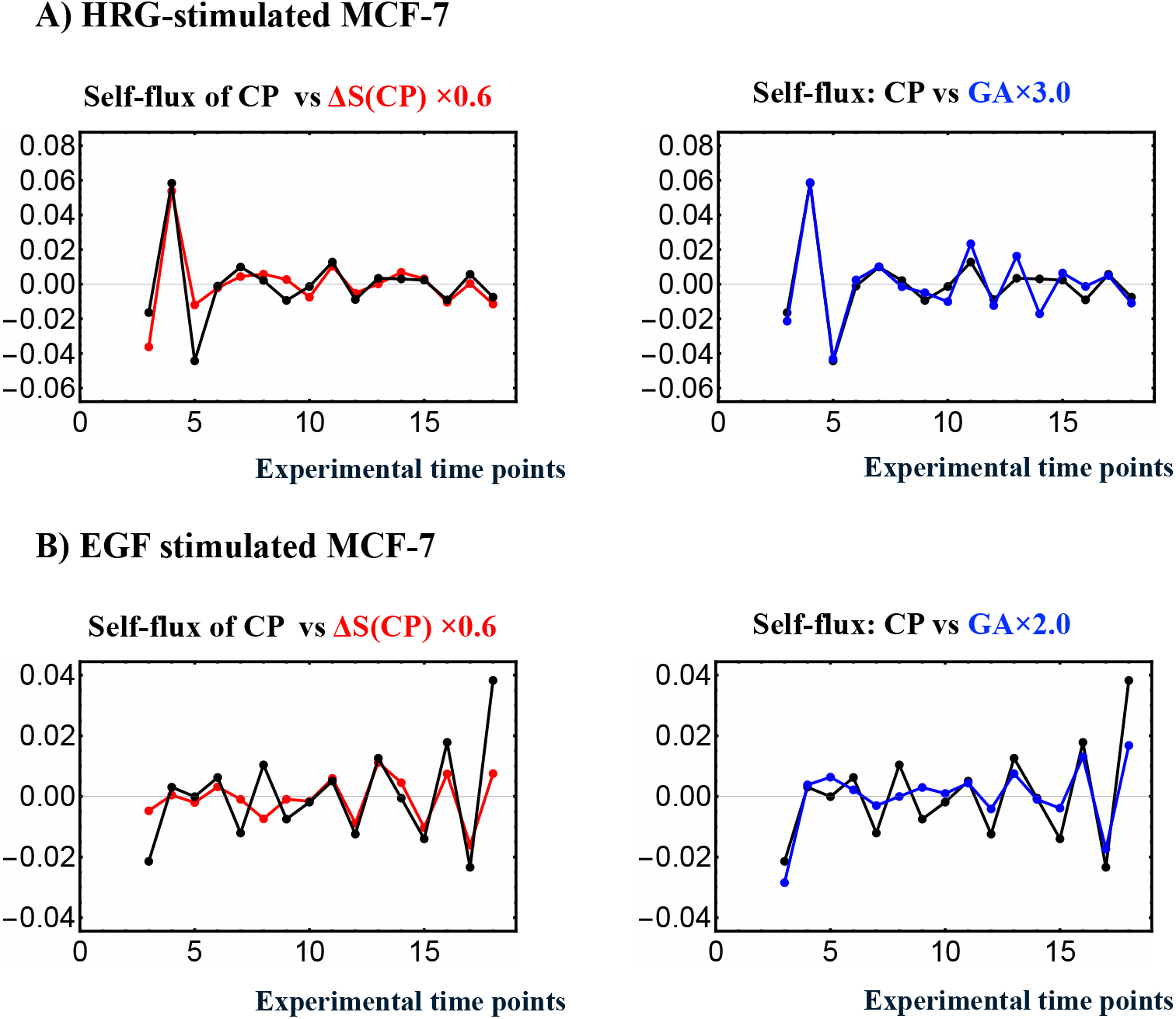
Concurrent occurrence of entropy-based thermodynamic synchronization and force-like dynamical synchronization. **A)** In HRG stimulation and **B)** EGF stimulation, the left panel shows the phase synchrony between self-flux (a dynamical system measure) and entropy change ΔS(CP) (a thermodynamic measure) for CP genes, while the right panel presents the self-flux of the CP and genome attractor (GA). The self-flux for the CP and GA is defined as the second difference from its overall temporal average value: -(*ε*(t_j+1_) - 2·*ε*(*t*_j_) + *ε*(t_j-1_)), where *ε*(*t*_j_) represents expression of the center of mass of CP genes or GA at *t* = *t*_j_. The negative sign arises from the correspondence with a harmonic oscillator when the system becomes linear. The change in CP entropy is given by its first difference: *S*(CP(*t*_j_)) - *S*(CP(*t*_j-1_)). Phase synchrony becomes apparent after the 10-min, with Δ*S*(CP) after 10-15 min and self-flux after 0-10-15 min. HRG stimulation, which induces a critical transition, shows stronger phase synchrony compared to EGF stimulation, where no critical transition occurs. In contrast, in the EGF case, the early-time CP formation-deformation cycle suggests that it leads to a weak phase synchrony among CP self-flux, its entropy change, and GA self-flux (see **Figure 8A**).

### 2) Autonomous Genome Computing as a Conceptual Framework

Our study demonstrates that net mutual information, incorporating higher-order nonlinear interactions, drives the critical transition that guides cell fate. This transition is governed by genomic thermodynamic phase synchronization and is mediated by CP genes functioning as rewritable chromatin memory. This framework grounds the concept of autonomous genome computing in measurable, mechanistic processes, where the genome integrates regulatory inputs and dynamically transitions between distinct functional states.

Recent advances suggest that genomic DNA is not merely a static blueprint but rather a dynamic, computationally capable system - a concept we term “genome computing.” In this framework, “computation” refers to the genome’s autonomous ability to integrate diverse regulatory inputs, store adaptive information, and transition between functional states in a context-dependent manner. This perspective unifies our understanding of the nonlinear and adaptive behaviors that regulate gene expression, drive cell differentiation, and govern cellular responses to external cues.

One key insight is that time-dependent transitions in genomic states can be modeled using kinetic equations with cubic nonlinearity, derived from symmetry arguments in thermodynamics[Takenaka et al., 2008; Nagahara and Yoshikawa, 2010]. This nonlinearity arises from changes in translational and conformational entropy within large DNA molecules interacting with their associated ionic environments [Nakai et al., 2005; Luckel et al., 2005; Tsuji et al., 2010; Nishio et al., 2021, 2022]. Experimentally observed chromatin reorganizations support this perspective, suggesting that genomic architecture is continuously tuned to explore and settle into energetically favorable configurations.

The symmetry characterized by cubic nonlinearity mirrors excitability phenomena observed in neuronal systems, as described by the FitzHugh-Nagumo [FitzHugh, 1955; Nagumo et al., 1962] and Hodgkin-Huxley [Hodgkin and Huxley, 1952] models. While these parallels remain conceptual, they underscore a shared principle: nonlinear systems, whether neural or genomic, can exhibit threshold-dependent switching between stable states.

As demonstrated in **Figure 2** and **section 2.6**, the CP region exhibits large-scale bimodal singular behavior based on *ln*(*nrmsf*) values. Consistent with this, recent evidence on bimodality clearly demonstrates that genome-sized DNA exhibits a bimodal free energy profile [Yoshikawa et al., 2000; Iwaki et al., 2003; Nakai et al., 2005; Mayama et al., 2007; Estévez-Torres et al., 2011; Yoshikawa et al., 2011; Nishio et al., 2020]. This bimodal symmetry inherently implies cubic nonlinearity in the reaction kinetics, which arises from the functional derivative of the free energy [Yoshikawa, 2002; Takenaka et al., 2008; Nagahara and Yoshikawa, 2010; Kanemura et al., 2018].

At the heart of these transitions lies the critical point (CP). While its precise molecular composition remains to be fully delineated, existing evidence supports that specific genomic regions consistently orchestrate large-scale regulatory shifts. As cells approach critical decision points such as lineage commitment, the CP emerges from bimodal states, toggling chromatin configurations between more and less active forms. Observations of pericentromere-associated domain (PAD) rearrangements and chromatin remodeling [Krigerts et al., 2021] further support the notion that the CP governs these critical transitions, effectively “choosing” among distinct genomic states.

These transitions can be conceptualized through an energy landscape perspective, where changes in translational and conformational entropy reshape the free energy landscape, favoring certain pathways over others. Analogies with Ginzburg-Landau theory [Ginzburg and Landau, 1965] highlight the importance of identifying order parameters (e.g., chromatin compaction or gene network states) and recognizing that critical transitions occur when specific thresholds are crossed. Our self-organized criticality (SOC) further extends this view: the genome, via the CP, naturally tunes itself to critical points, maintaining a balance between order and flexibility. Acting as rewritable chromatin memory, the CP plays a role akin to Maxwell’s demon, selectively promoting transitions that reduce uncertainty and drive the system toward coherent and functionally relevant states.

Importantly, this notion of “genome computing” does not imply that the genome functions like a digital processor with discrete inputs and outputs. Instead, genome computing is characterized by autonomous, decentralized super-information processing. It reflects a continuous, adaptive reshaping of genomic states, where “memory” is encoded in the genome’s ability to revisit specific conformations, and “logic” emerges from network interactions that drive transitions toward or away from stable regulatory configurations.

In conjunction with the genomic-thermodynamic mechanism, understanding how the CP functions as a central process hub to orchestrate the complex spatiotemporal self-organization of the genome will elucidate fundamental principles governing cell fate, development, and stress responses. This knowledge could ultimately inspire innovative approaches in regenerative medicine, cancer therapy, and synthetic biology.

### 3) Future Prospects for Genome Intelligence

At a more theoretical level, the concept of Genome Intelligence (GI) builds upon the computational principles of genome computing to describe the genome’s capacity for integrated, adaptive, and memory-like behavior. Here, intelligence refers to a minimal form of integrated responsiveness, in which the genome not only reacts to environmental stimuli but also incorporates these inputs into its regulatory framework, enabling adaptation and future decision-making.

Integrated Information Theory (IIT), originally developed to explain consciousness in neural systems, offers a useful framework for understanding intelligence in biological systems more broadly. The fundamental challenge addressed by IIT is how to integrate external informative stimuli into the structure of an “intelligent agent,” leading to the emergence of a memory of the stimulus. This stands in contrast to non-intelligent sensors (e.g., a photodiode), whose structure remains unchanged by interactions with stimuli.

As demonstrated by Niizato et al. (2024), IIT extends beyond neural systems to encompass molecular and cellular systems, where complex interactions give rise to irreducible wholes, systems that cannot be decomposed into independent parts without a loss of function. When applied to the genome, GI emerges from the integration of external signals into chromatin configurations, regulatory networks, and gene expression states. This capacity for integration, persistence, and reconfiguration distinguishes the genome as an autonomous, intelligent system in a minimal yet fundamental sense.

It is important to clarify that referring to GI does not imply a claim that the genome possesses intelligence in the fully defined, cognitive sense. The term “intelligence” derives from the Latin *intus* + *legere*, meaning “to read within” - to uncover hidden or implicit knowledge. While the genome lacks intentional insight, referring to Genome Intelligence emphasizes its ability to track past experiences, as demonstrated by epigenetic memory (Elsherbiny and Dobreva, 2021).

Ultimately, GI reframes our understanding of the genome as a dynamic, adaptive, and integrative system. In our view, GI highlights the consilience between computational and thermodynamic principles and biological mechanisms, offering a foundation for exploring how genomes encode past experiences, adapt to environmental changes, and guide cellular behavior. This perspective has profound implications for fields such as regenerative medicine and synthetic biology, where leveraging the intelligence-like properties of the genome may transform how we design and manipulate living systems.

## 4. Conclusion

Our study demonstrates that genomic regulation can be understood through the lens of open stochastic thermodynamics. Rather than existing at thermodynamic equilibrium, gene expression operates far from equilibrium, with continuous energy dissipation enabling dynamic regulation, adaptability, and responsiveness to external cues. By examining the roles of critical point (CP) genes and the whole expression system (WES) in MCF-7 breast cancer cells,we establish a thermodynamic framework that explains how biological systems autonomously maintain coherence, stability, and flexibility in their gene expression profiles.

Key insights from our findings include the thermodynamic phase synchronization of CP genes with the genome, the distinct roles of the CP under different stimuli, and the presence of positive higher-order mutual information. The latter highlights nonlinear interdependencies, enhancing both synergy (emergent collective information) and redundancy (overlapping information) beyond the pairwise-interaction framework among genes.

Under epidermal growth factor (EGF) stimulation, the CP remains passive, preserving stability without transitioning into a new state. In contrast, under heregulin (HRG) stimulation, the CP “reads” and redistributes information through rewritable chromatin memory, exhibiting a Maxwell’s demon-like function to orchestrate a global shift in gene expression. In other words, the genome “learns” from environmental signals, processes this information, and then reprograms its overall state - a phenomenon we describe as “Genome Intelligence (GI)”.

On a broader level, these computational dynamics support the concept of GI, where the genome exhibits emergent properties such as the ability to discriminate between environmental states, integrate signals into its structural framework, and adaptively guide future regulatory transitions. This emergent intelligence reflects the genome’s capacity to store “memory” of past states and utilize this information to navigate complex decision-making landscapes. These insights bridge classical molecular genetics with non-equilibrium thermodynamic and computational principles, providing a holistic view of genome regulation.

Our findings open new avenues for understanding disease progression, guiding regenerative medicine, and informing innovative cancer therapies. For example, the CP’s ability to dynamically “compute” and integrate signals highlights opportunities for targeting critical regulatory hubs in cancer treatment or leveraging genomic intelligence in synthetic biology to design adaptive, programmable cellular systems.

From a global perspective, integrating genome computing with principles of intelligence inspires the development of machine learning tools modeled on cell fate transition dynamics. Physics-Informed Neural Networks (PINNs), as described by Cai et al. (2021), offer a promising framework for incorporating physical principles into neural network architectures to replicate the behavior of biological systems. Similarly, Hopfield networks, as noted by Krotov [Krotov, 2023], demonstrate how network correlations can encode emergent behavior, much like genomic systems encode adaptive regulatory transitions. By applying these computational principles as hyper-parallel, autonomous, decentralized systems, we can enhance both biological understanding and machine learning, paving the way for models that seamlessly integrate thermodynamic, computational, and biological insights.

## 5. Material

Microarray data for the activation of ErbB receptor ligands in human breast cancer MCF-7 cells by EGF and HRG were obtained from the Gene Expression Omnibus (GEO) ID: GSE13009 (N = 22,277 mRNAs; for experimental details see [Saeki et al., 2009]) and measured at 18 time points: t_1_ = 0, t_2_ = 10, 15, 20, 30, 45, 60, 90 min, 2, 3, 4, 6, 8, 12, 24, 36, 48, t_18_ = 72 h. Each condition includes two replicates (rep 1 and rep 2); the analyses presented in this report are based on rep 1 for both EGF and HRG, while the results from rep 2 of HRG stimulation are provided in the **Supplemental File**. The robust multichip average (RMA) was used to normalize expression data for background adjustment and to reduce false positives [Bolstad et al., 2003; Irizarry et al., 2003; McClintick and Edenberg, 2006]. Refer to RNA-Seq data processing details in **section 2.1**‥

## Abbreviations

CM: center of mass
CP: critical point
CSB: coherent stochastic behavior
cSOC: classical self-organized criticality
EGF: epidermal growth factor
GA: genome attractor
GI: genome intelligence
HRG: heregulin
IIT: integrated information theory
ITA: information thermodynamics analysis
MI: mutual information
min: minutes
nrmsf: normalized root mean square fluctuation
OMG genes: oscillating-mode genes
PCA: principal component analysis
PC1: first principal component
PC2: second principal component
PES: peripheral expression system
WES: whole expression system.

## 6. Appendix Mathematical description of Chromatin Entropy

As illustrated in **Figure 7**, we transform the gene expression distribution into the normalized root mean square fluctuation (*nrmsf*) probability distribution—an entropy decomposition approach analogous to the logical/physical state framework described by Maroney [Maroney, 2009]. To provide a mathematical perspective, we describe chromatin entropy by showing how the Shannon entropy of the Whole Expression System (WES), defined in equation (2), decomposed into “logical states”, each corresponding to *nrmsf* bins as follows:

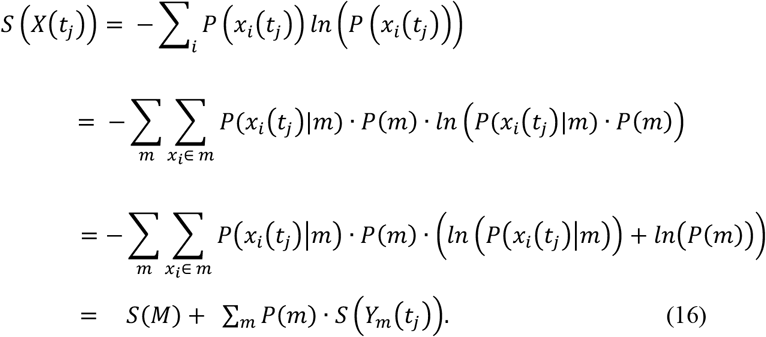

Here, the probability *P*(*x*_i_(*t*_j_)) of gene *x*_i_ being expressed at time *t*_j_ can be factored into 1) the probability of selecting a logical state *m, P*(*m*), and 2) the conditional probability of the physical state *x*_i_(*t*_j_) within that logical state, *P*(*x*_i_(*t*_j_)|m), such that *P*(*x*_i_(*t*_j_)) = *P*(*x*_i_(*t*_j_)|m)· *P*(*m*). The sum of conditional probabilities within each logical state satisfies 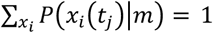 at time *t*_j_.

The entropy of the whole expression system, *S*(WES), is thus expanded into two components: *S*(*M*), the entropy associated with the distribution of logical states, and *S*(*Y*_m_(*t*_j_)), which represents the entropy of the physical states (gene expression levels) within the *m*^th^ logical state:

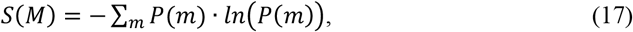

and

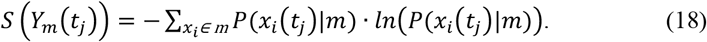

## Author Contribution

MT: Conceptualization (lead); Formal analysis and Software; Methodology; Writing – original draft; Writing – review & editing

KY: Conceptualization (supporting); Writing – review & editing

AG: Conceptualization (supporting); Writing – review & editing

## Acknowledgments

MT thanks Prof. Paul Brazhnik for his insightful discussions and the SEIKO Life Science Laboratory, Osaka, Japan, for its invaluable support during this research. He is also grateful to his family (Ms. Takako Tsuchiya; Drs. Kimiko and Kazumi Tsuchiya; and Dr. Harry Taylor) for their constant encouragement. Special thanks go to Dr. Daisaku Ikeda for his lifelong inspiration, and to Ms. Satoko Takeda and Ms. Hisami Spector for their kind and heartfelt support.

## Supplemental File

Our information-thermodynamics analysis (ITA) demonstrates that the CP functions as a Maxwell’s demon in genome regulation. Two additional supporting results, labeled **I** and **II**, are provided in this Supplemental File.

## I) Independent Replicate Result (Replicate 2) for HRG-Stimulated MCF-7 Cells

Based on our information-thermodynamics analysis (ITA), we present results from replicate 2 of the MCF-7 gene expression dataset under HRG stimulation [Saeki et al., 2009], available in the Gene Expression Omnibus (GEO) under accession ID **GSE13009**. The figure numbers correspond to those used in the main text. The results, <u>consistent with those from</u>**<u>Replicate 1</u>**, reveal CP–PES phase synchronization accompanied by the activation of the Maxwell’s demon function, as detailed below:

**Figure S1.**
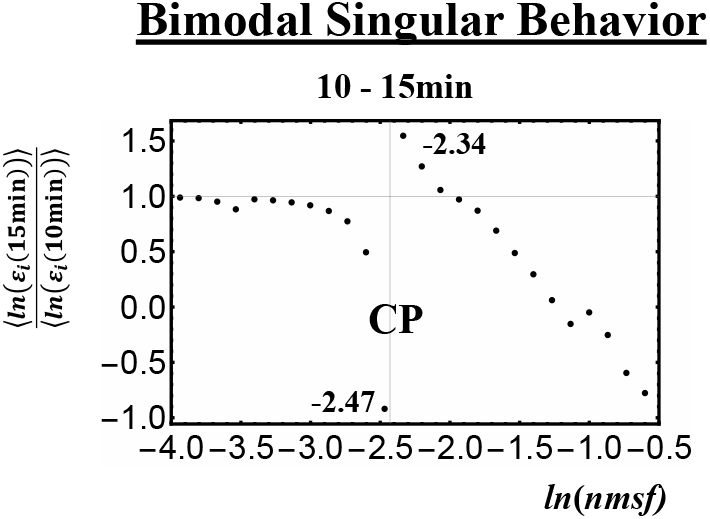
Identification of the Critical Point (CP) Region. The CP region exhibits bimodal singular behavior at 10-15 min. The peak range spans from -2.47 to -2.34 in *ln*(*nrmsf*). The CP region is defined as -2.5 < *ln*(*nrmsf*) < -2.3, encompassing 3,598 genes (see the **Figure 2** caption in the main text for details).

**Figure S2.**
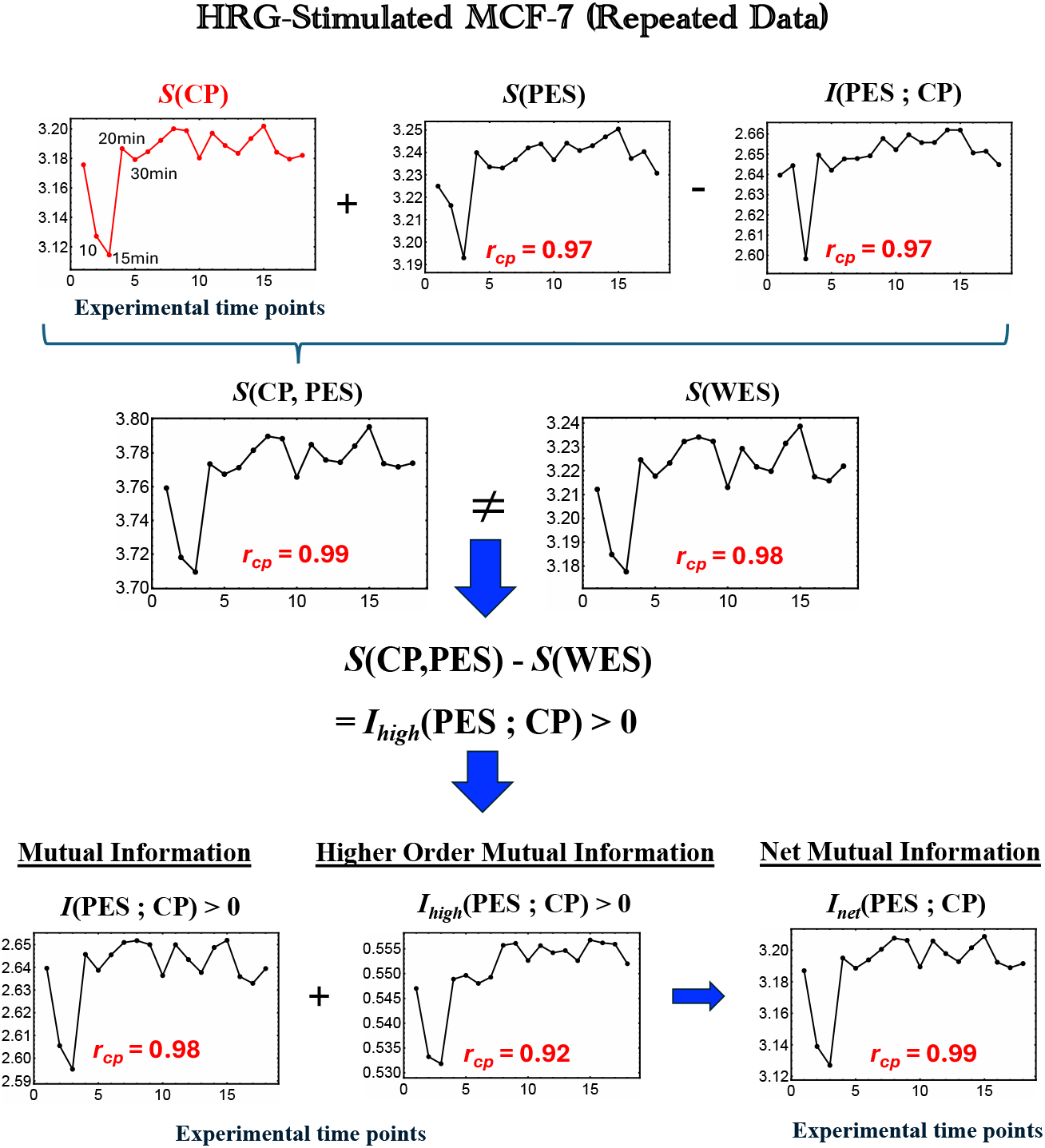
Thermodynamic Phase Synchronization and Higher-Order Mutual Information in HRG-Stimulated MCF-7 Cells (Replicate 2). The procedure follows that described in Figure 5 in the main text.

1. **CP-PES synchronization is also evident in Replicate 2** for HRG-stimulated MCF-7 cells, as demonstrated by an almost perfect temporal Pearson correlation between the entropy of the critical point, **S(CP), and** the net mutual information, **I**_**net**_**(CP;PES)**.
2. A positive higher-order mutual information **I**_**high**_**(CP;PES) > 0** indicates cooperative thermodynamic behavior between the critical point (CP) and the peripheral expression system (PES). The net mutual information, **I**_**net**_**(CP;PES)** exhibits a pulse-like pattern, <u>coinciding with the timing of a critical transition in the genome engine mechanism</u> [Tsuchiya et al., 2020, 2022, 2023a]. The following figure visualizes this pulse-like behavior in **I**_**net**_**(CP;PES)**, revealing a <u>three-step process</u> involving a Maxwell’s demon-like action that rewrites chromatin as dynamic memory.

**Figure S3.**
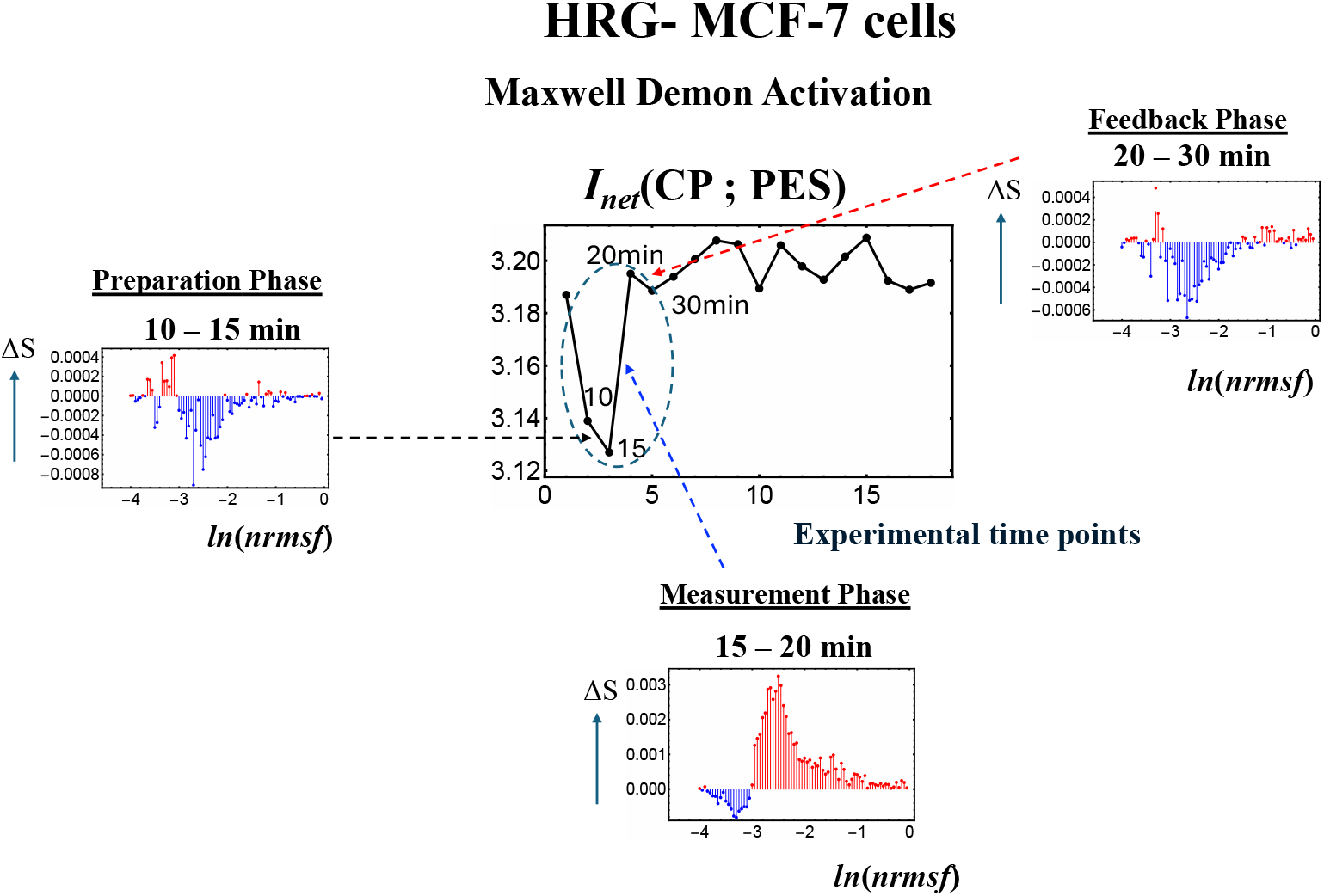
Maxwell’s Demon Phases in HRG-Stimulated MCF-7 Cells (Replicate 2). The net mutual information, *I*_net_(CP; PES), under HRG stimulation reveals all three operational phases of a Maxwell’s demon: **preparation** (10-15 min), **measurement** (15–20 min), and **feedback associated with the critical transition** (20-30 min), accompanied by coherent chromatin remodeling dynamics. (see **Figure 9** caption in the main text for details). <u>This result supports the findings from Replicate 1 in our study</u>.

### II) DMSO-Stimulated HL-60 Human Leukemia Cells

Using GEO ID: **GSE14500** (N = 12,625 mRNAs; details in [Huang et al., 2005]) **across** 13 time points: t_1_ = 0, t_2_ = 2, 4, 8, 12, 18, 24, 48, 72, 96, 120, 144, t_13_ = 168 h, ITA reveals pulse-like behavior in ***I***_**net**_**(CP; PES)**, exhibiting a three-step process involving a Maxwell’s demon-like action that rewrites chromatin as dynamic memory. This further demonstrates that pluse-like net mutual information **reflects the CP’s function as a Maxwell’s demon in genome regulation**.

**Figure S4.**
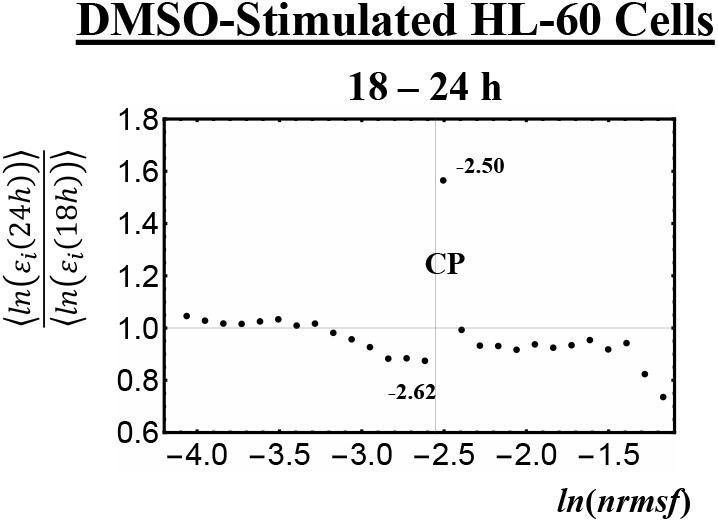
Identification of the Critical Point (CP) Region. The CP region exhibits bimodal singular behavior at 18 -24 h. The peak range spans from -2.62 to -2.50 in *ln*(*nrmsf*). The CP region is defined as -2.65 < *ln*(*nrmsf*) < -2.45, encompassing 1,719 genes (see the **Figure 2** caption in the main text for details).

**Figure S5.**
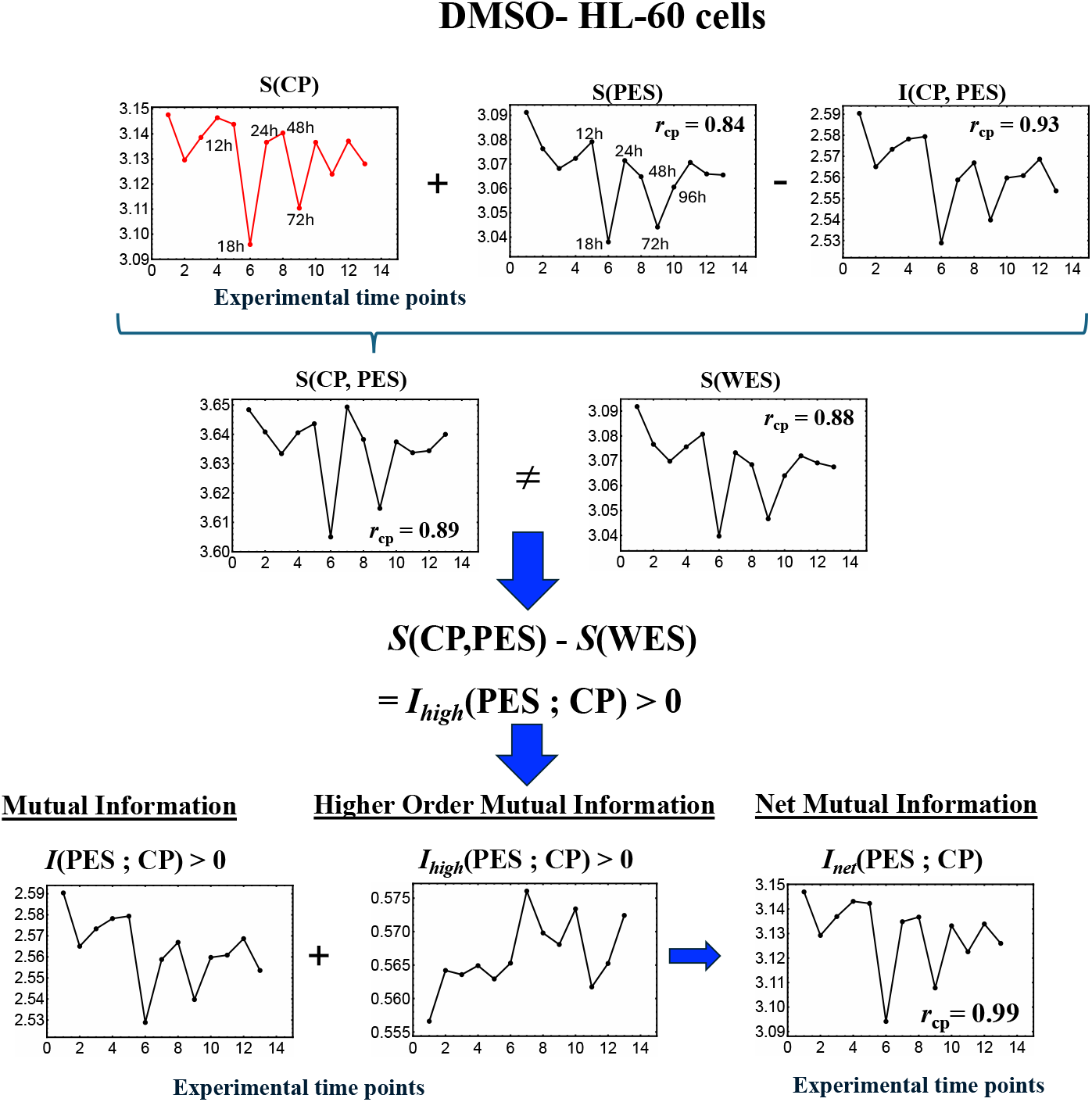
Thermodynamic Phase Synchronization and Higher-Order Mutual Information in DMSO-Stimulated HL-60 Cells. **<u>Thermodynamic phase synchronization and positive higher-order mutual information are also evident</u>**<u>in</u>**<u>DMSO-stimulated HL-60 cells</u>**, indicating cooperative behavior between the critical point (CP) and the peripheral expression system (PES).

**Figure S6** visualizes the pulse-like behavior in ***I***_**net**_**(CP; PES)**, revealing a three-step process involving a Maxwell’s demon-like action that rewrites chromatin as dynamic memory. The net mutual information, ***I***_**net**_**(CP; PES)**, exhibits a pulse-like pattern at 12-18 h (**preparation**), 18-24 h (**measurement**), and 48-72 h (**feedback**), accompanied by **coherent chromatin remodeling.** The critical transition corresponding to feedback occurs at 48 h.

**Note:** The genome engine mechanism exhibits global expression changes (genome avalanche) at 12-18 h and 18-24 h, corresponding to the pulse-like changes in *I*_net_(CP; PES) observed in **Figure S6**. Genome engine switching occurs at 18 h, indicating a

**Figure S6.**
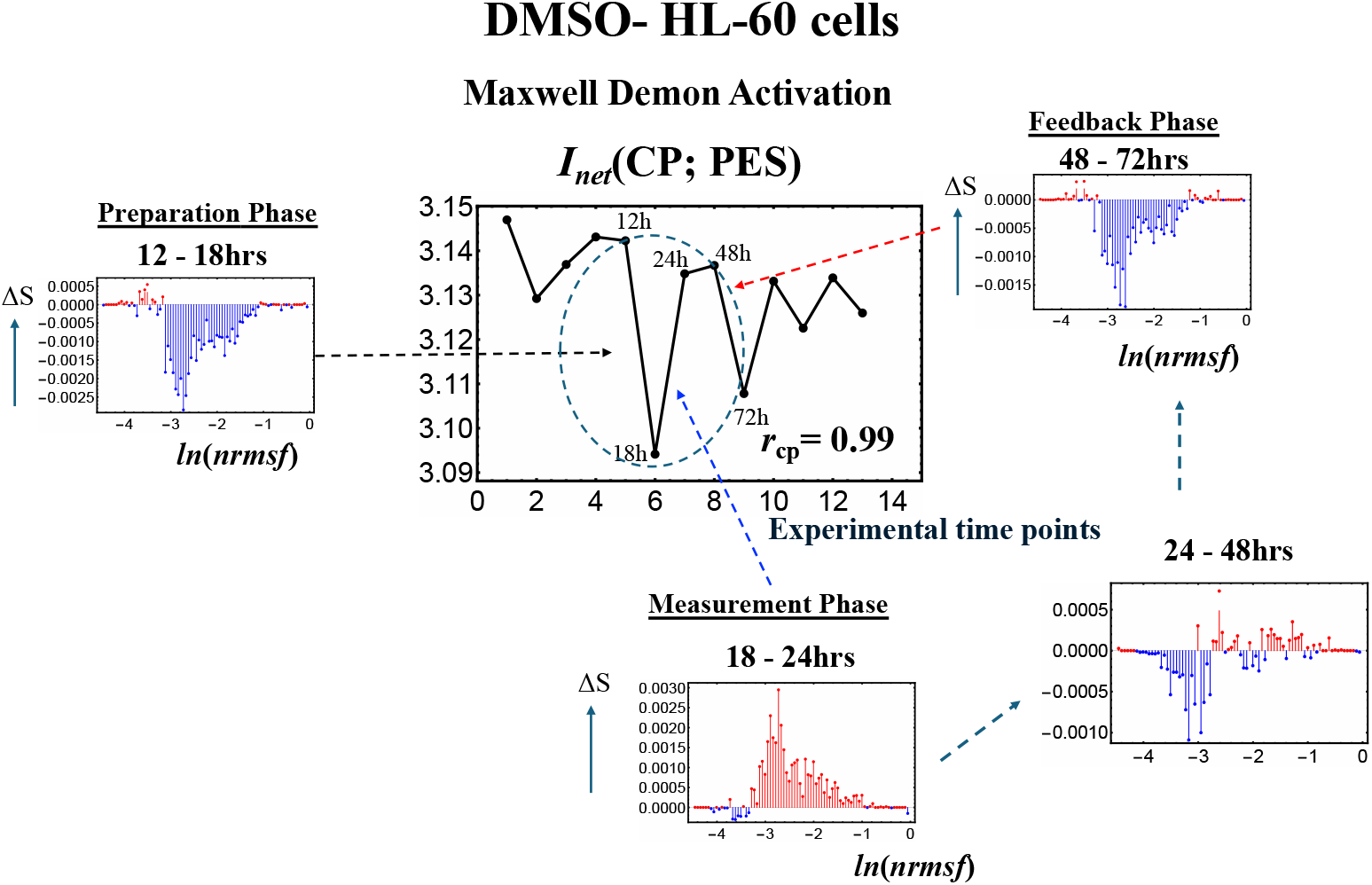
Maxwell demon function as rewritable chromain memory in DMSO stimulated HL-60 Cells. The net mutual information **I**_**net**_**(CP;PES)** reveals a three-phase process - **preparation (12-18 h), measurement (18-24 h)**, and **feedback (48-72 h)** - characteristic of a dynamic Maxwell’s demon-like mechanism. This process reflects the function of rewritable chromatin memory during the critical transition induced by DMSO stimulation.

## Notes

Conflict of Interest: The authors declare no conflict of interest.

### Competing Interest Statement

The authors have declared no competing interest.

### Summary of Updates

We have substantially revised the manuscript to improve clarity and reinforce the core finding: Maxwell's demon activation under genomic thermodynamic phase synchronization. Key updates in the main text include: 1.A new Figure 1 outlining the step-by-step information-thermodynamics workflow for both microarray and RNA-seq data. 2.A new Figure 10 showing power-law scaling of chromocentre distribution in MCF-7 cells under HRG stimulation. 3.Updated Figures 7 and 9. 4.Expanded all figure captions with clearer, more informative annotations. 5.New experimental evidence in section 2.7 demonstrating that ln(nrmsf)-based gene-expression variability reflects chromatin remodeling dynamics. 6.A Supplemental File (attached after the References section) further supporting CP-driven Maxwell's demon activity in an independent replicate (HRG-stimulated MCF-7 cells) and in a different cancer cell type: DMSO-stimulated HL-60 cells.

